# Point-of-care analyte quantification and digital readout via lysate-based cell-free biosensors interfaced with personal glucose monitors

**DOI:** 10.1101/2021.06.22.449464

**Authors:** Yan Zhang, Paige L. Steppe, Maxwell W. Kazman, Mark P. Styczynski

**Author notes:** These authors contributed equally to this work (Paige L. Steppe, Maxwell W. Kazman).

## Abstract

Field-deployable diagnostics based on cell-free systems have advanced greatly, but on-site quantification of target analytes remains a challenge. Here we demonstrate that *Escherichia coli* lysate-based cell-free biosensors coupled to a personal glucose monitor (PGM) can enable on-site analyte quantification, with the potential for straightforward reconfigurability to diverse types of analytes. We show that analyte-responsive regulators of transcription and translation can modulate production of the reporter enzyme β-galactosidase, which in turn converts lactose into glucose for PGM quantification. Because glycolysis is active in the lysate and would readily deplete converted glucose, we decoupled enzyme production and glucose conversion to increase endpoint signal output. This lysate metabolism did, however, allow for one-pot removal of glucose present in complex samples (like human serum) without confounding target quantification. Taken together, we show that integrating lysate-based cell-free biosensors with PGMs enables accessible target detection and quantification at the point of need.

**Table of Contents Graphic:** 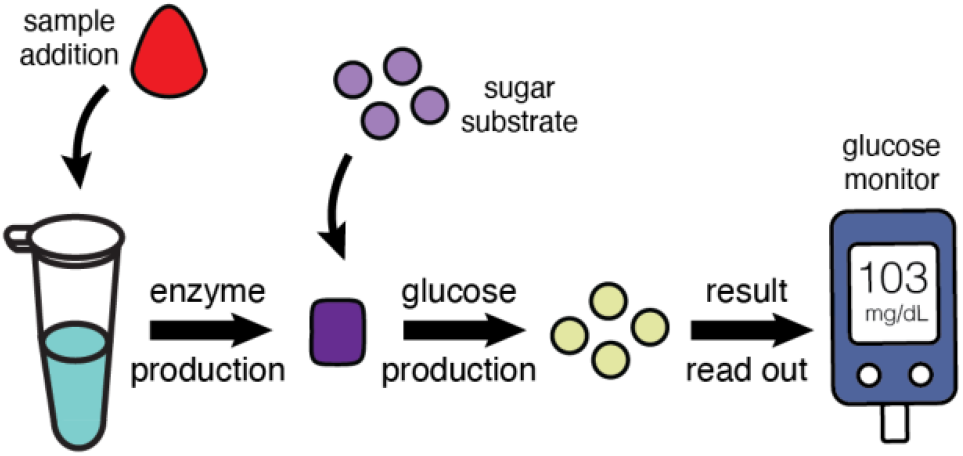

## Introduction

Cell-free expression (CFE)-based biosensors hold great potential for onsite measurement of target analytes. A freeze-dried pellet consisting of the core protein expression machinery—as is the result from preparing bacterial lysates for cell-free expression—plus plasmid DNA coding for biosensor and reporter output can be sufficient to let a worker in the field perform sample analysis at the sampling site. This strategy avoids the complicated logistics that are otherwise required to bring samples to appropriately equipped and staff laboratories. The extremely low sample volume, operator, and infrastructure requirements for CFE-based sensors make this platform particularly promising for development of low-cost and easy-to-use diagnostic tools for applications ranging from healthcare screenings to environmental surveillance^1^.

Despite the fact that applications of CFE systems for biosensing have expanded in recent years, rapid and reliable analyte quantification in CFE reactions at the point of care remains a challenge. Most of the current on-site CFE detection strategies use enzymatic reporters to generate visible color pigment for either a binary, yes-or-no result readout^2, 3^ or a semi-quantitative measurement of target concentration using transient color changes^4^. Others have used custom-built, portable electronic devices in place of bulky plate readers for result interpretation and quantification^2, 3, 5^. While these efforts represent promising strides toward quantitative analyte measurement at the sampling site, few of these approaches or devices can match the simplicity, quantification, and digital readout offered by a personal glucose monitor (PGM).

Since the product’s commercialization in the 1970s, personal glucose monitors have been through decades of refinement and matured into a robust and easily accessible technology that many patients use daily and almost anywhere^6^. As a result, there has been significant research effort to engineer biosensors that can be read by a glucose monitor^7–15^ rather than trying to engineer entirely new quantification devices that match the PGM’s portability and reliability. Recent efforts have even successfully interfaced glucose monitors with cell-free system-based biosensors, though they were subject to significant limitations. The first-ever reported use of cell-free systems with a glucose monitor was limited to using reagent complementation strategies^8^, such that it was only used to measure analytes that were required components of the cell-free reaction. A more recent report demonstrated detection of the presence and absence of targets (nucleic acid sequences) but did not aim for analyte quantification^7^. In addition, the demonstration used an expensive, purified protein expression system (PURExpress, at almost $10 per sample) and required a separate, overnight enzymatic conversion step to remove confounding glucose that is present in the sample. The glucose removal step that was used required the user to know in advance the sample glucose levels and then add sample-specific volumes of reagents to clear native glucose from samples without causing unwanted degradation of the glucose generated by the sensing reaction. All of these requirements decrease that approach’s potential viability for field-deployable applications. While that report was a significant step forward, the use of a PGM for qualitative, presence/absence diagnosis rather than quantitative measurement meant the strategy did not fully exploit one of the most critical and impactful capabilities offered by PGMs. Even more importantly, for most clinically relevant biomarkers for conditions other than infectious disease, it is the biomarkers’ concentration—not merely their presence or absence—that is the criterion for diagnosis^4, 16–18^. For these target analytes, quantification at the point of need is critical and digital readout enables straightforward result interpretation.

Here, we aim to expand the repertoire of glucose monitor-mediated analyte detection to include quantification of diverse sets of analytes via lysate-based CFE systems. We first showed that a genetic circuit constitutively expressing the enzyme β-galactosidase (LacZ) in the CFE lysate reaction successfully allows LacZ conversion of lactose to glucose and yields measurable PGM outputs. We then demonstrated analyte-modulated LacZ production and glucose conversion via different biosensing circuits. We successfully used a zinc-responsive transcription factor to generate dose-dependent expression of LacZ to identify zinc deficiency in a human serum matrix at clinically relevant concentrations. We further showed that the same detection strategy could be used to detect and quantify nucleic acid biomarkers from pathogenic *E. coli* by merely substituting the zinc transcriptional control elements with RNA regulatory elements (toehold switches). In developing these diagnostic sensors, we found that the metabolic pathways active in lysate-based CFE systems^19–23^ readily deplete glucose in reactions. As a result, we decoupled LacZ production from glucose conversion and capitalized on this lysate metabolism for one-pot removal of glucose that may be initially present in a complex sample environment (like human serum or other biofluids) without customizing reagent volumes to individual samples, thereby eliminating a separate processing step to remove endogenous glucose required in current PGM-mediated analyte quantification methods^7, 14, 15^. Taken together, our work showcases a broadly applicable and modular strategy for rapid and reliable quantification of target analytes at the point of need, expanding the repertoire of PGM-mediated biomarker detection with biosensors expressed in lysate-based CFE systems.

## Results and Discussion

### Lysate-based CFE reaction producing LacZ enzyme can convert lactose to measurable glucose

The first step toward PGM-mediated analyte quantification using lysate-based CFE reactions was to confirm that the CFE reagents are compatible with commercial PGMs. To verify reagent compatibility and glucose stability, we incubated D-(+)-glucose (hereafter referred to as glucose) at concentrations of 0-25 mM in CFE reactions to span the full PGM detection range and tracked their respective value readouts over time. Compared to glucose standards prepared in water, we observed a slight decrease in signal output for immediate measurement of the same glucose concentration in the CFE matrix (**Figure 1A**). We also found significant glucose consumption in CFE reactions over time (**Figure 1A**). This finding was perhaps unsurprising, as previous reports have observed and characterized significant endogenous glycolytic metabolic activity in CFE lysates^19–23^; we attributed the loss in signal output over time to enzyme-catalyzed conversion of glucose to glucose-6-phosphate due to residual glycolytic activity in the lysate.

**Figure 1:**
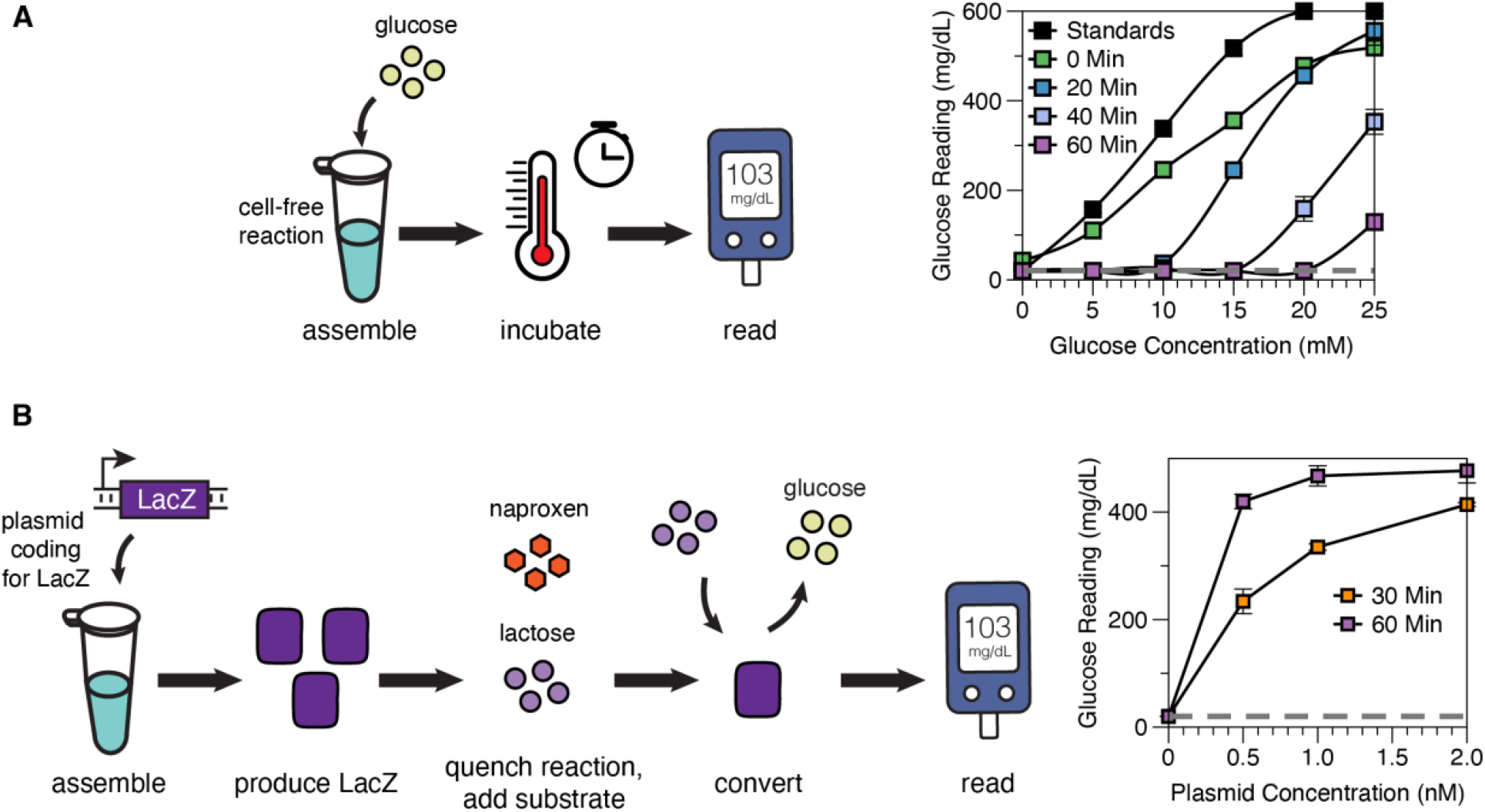
Characterization of glucose production and depletion in CFE reactions. **(A)** Verification of CFE compatibility with PGM and time-course measurement of glucose depletion in reactions. A slight decrease in PGM output was observed for the same concentration of glucose in the CFE matrix and no incubation time compared to glucose in a water solution (“Standards”). Rapid depletion of glucose signal was observed in all CFE reactions over time. Error bars represent the standard deviation of cell-free reaction triplicates. Dashed gray line represents PGM’s lowest reading threshold, 20 mg/dL. **(B)** Decoupling enzyme production and glucose conversion in CFE reactions enabled dose-dependent PGM signal output. CFE reactions containing varying concentrations of plasmid constitutively expressing LacZ were incubated for 30 to 60 minutes before each reaction was quenched by naproxen-lactose mix to shift the reaction from enzyme production to glucose conversion. Plasmid concentration-modulated glucose production was detected using the PGM after 15 minutes of incubation. Error bars represent the standard deviation of cell-free reaction triplicates. Dashed gray line represents PGM’s lowest reading threshold, 20 mg/dL.

Endogenous glycolytic activity in crude *E. coli* lysate could pose serious problems for CFE-mediated analyte quantification using PGMs, since glucose molecules generated by the reporter enzyme in the CFE reaction would be readily depleted, and thus desired signal would be lost. To address this issue, we chose to decouple reporter enzyme production from enzyme-catalyzed glucose production. To assess how fast glucose can be produced for detection by the PGM, we added a plasmid for constitutive expression of the enzyme LacZ to the CFE reaction for 30 to 60 minutes. Following this incubation, a mixture consisting of naproxen and lactose was added to the CFE reaction to terminate transcription in the CFE system (via naproxen), slow down lysate metabolism (via naproxen), and start glucose production via LacZ conversion of lactose to glucose. Different concentrations of plasmid that constitutively express LacZ were used as a testbed model for our eventual goal of LacZ expression that increases based on the amount of analyte present. After 15 minutes of incubation, we observed plasmid dose-modulated glucose signal production on the PGM (**Figure 1B**). Naproxen was used here due to its effectiveness at inhibiting CFE reactions without impairing LacZ activity^4^ (**Figure S1A-D**). We anticipate that other small molecule inhibitors added at high concentrations could also be capable of halting the CFE reaction, but we chose naproxen here due to its minimal effects on LacZ-mediated glucose production, its inhibition of endogenous glucose depletion (**Figure S1E**), and our previous experience using naproxen with cell-free biosensors^4^.

### Repurposing a PGM to quantify micronutrients in human serum

After successfully verifying plasmid dose-dependent glucose production in CFE systems, we tested whether small molecule inducers could modulate dose-dependent glucose readings on PGMs. We chose zinc as our target analyte for PGM-mediated quantification due to its global health relevance (zinc deficiency is responsible for the deaths of 100,000 children under the age of five worldwide every year)^24, 25^ and our group’s previous experience in developing a semi-quantitative zinc biosensor^4^. The zinc sensor used here constitutively expresses (from the promoter P_T7_) a transcription factor ZntR, which in turn controls the expression of LacZ based on the concentration of zinc. Zinc binding activates ZntR, which turns on expression from its cognate promoter P_zntA_ for LacZ production (**Figure 2A**).

**Figure 2:**
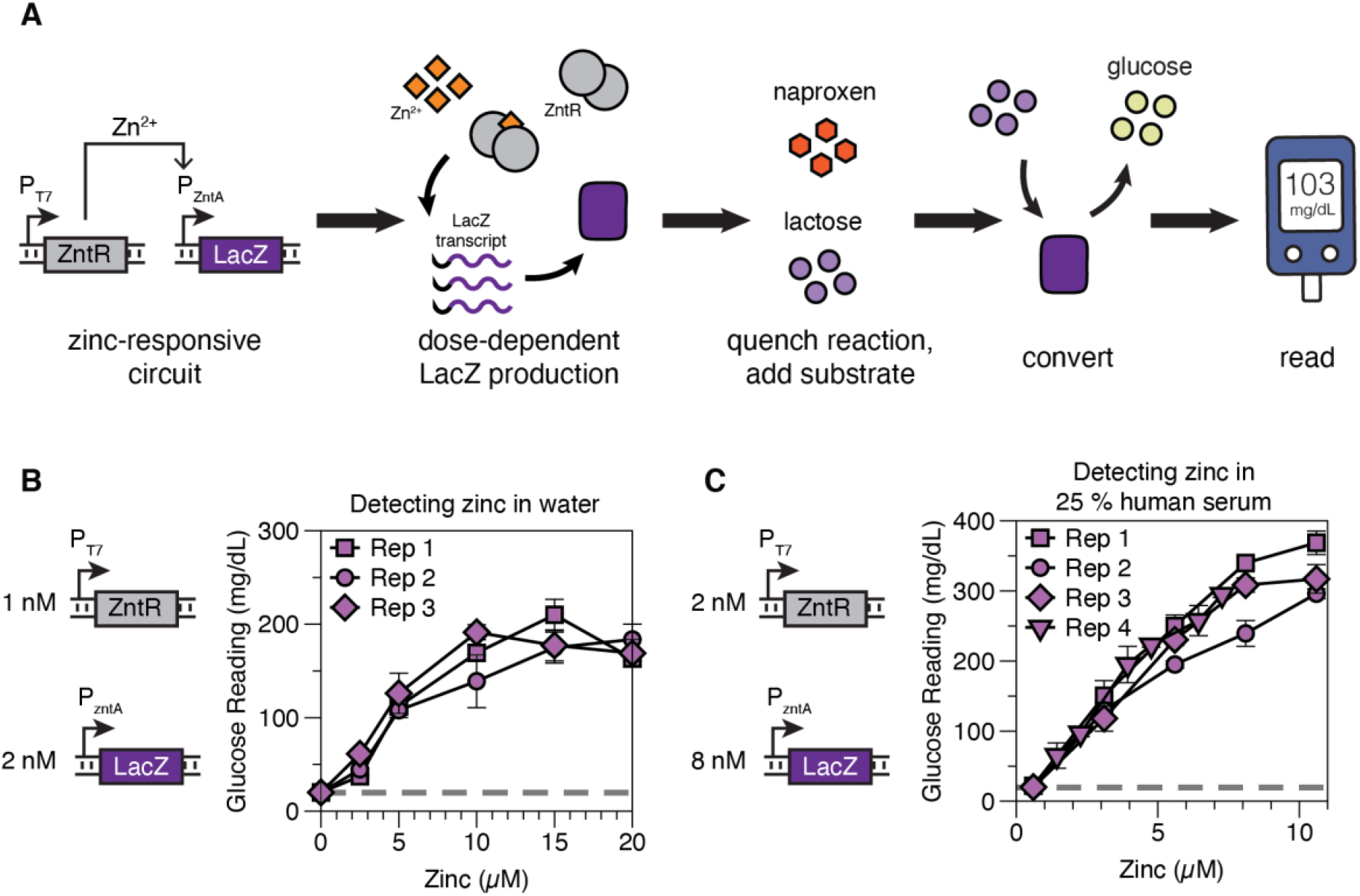
Application of PGM-mediated quantification of zinc in CFE reactions. **(A)** Schematic of zinc-modulated glucose production and PGM-mediated target quantification in CFE reaction. Zinc modulates LacZ production by binding to the constitutively expressed transcription factor ZntR, thereby activating transcription from the ZntR-responsive promoter P_zntA_. Following 45 min of LacZ production, a mixture of the naproxen-lactose solution was added to quench the CFE reaction and to start lactose conversion for 15 min. The converted glucose was then read on the PGM for target analyte quantification. **(B)** Dose-dependent glucose production in CFE reaction with zinc in a water matrix. The same experiment was replicated on different days to verify consistency in glucose output. Replicates (Rep) represent independently assembled reactions and error bars represent the standard deviation of cell-free reaction triplicates in each replicate. Dashed gray line represents PGM’s lowest reading threshold, 20 mg/dL. **(C)** Dose-dependent glucose production in CFE reaction with zinc in 25% pooled human serum. X-axis zinc concentrations reflect the total zinc in the reaction after accounting for the remaining zinc in chelated serum (Figure S2D). The same experiment was replicated on different days and with an independently assembled reaction to verify consistency in glucose output. Error bars represent the standard deviation of cell-free reaction triplicates in each replicate. Dashed gray line represents PGM’s lowest reading threshold, 20 mg/dL.

Because the human physiologically relevant zinc concentration spans from 2 to 20 μM^4, 24^, we first tested for zinc-modulated glucose production in a water matrix across this range (**Figure 2B**). Using the same strategy to decouple analyte detection from glucose conversion, we incubated CFE reactions for 45 minutes for LacZ production before quenching reactions with naproxen-lactose mix and incubating for another 15 minutes for glucose production. Although we have demonstrated that 30 min of reaction can generate enough LacZ to produce detectable glucose signals on the PGM, we extended the reaction to 45 min to account for the lag time associated with the expression of sufficient zinc-responsive transcription factor to enable expression of LacZ, an additional step not present in our initial experiments from Figure 1. In just 1 hour of total assay time (including glucose production), we observed a linear increase in glucose readout over a range spanning 0-10 μM zinc, above which the response of zinc sensor starts to saturate at increasing zinc concentrations^4^. Further, we observed consistent glucose production across different reactions assembled on different days, demonstrating that PGM-mediated analyte quantification could be a reliable method for daily monitoring of micronutrient status.

We then focused on the linear response range of zinc concentrations, re-optimized plasmid concentrations for increased signal, and tested the compatibility of our approach with human serum samples (**Figure 2C**). Because zinc is endogenously present in serum and the samples were from otherwise healthy volunteers, the baseline level of zinc in the pooled serum sample would prevent us from assessing the assay’s ability to detect deficient zinc levels. To address this issue, we first removed endogenous zinc from the pooled serum samples via chelation (see **Supplemental Method** and **Figure S2D**) and then spiked different concentrations of zinc back into the serum. In CFE reactions containing 25% pooled human serum, we observed a consistent dose-dependent glucose signal readout over a range spanning 0-10 μM zinc across different days (**Figure 2C**), reflecting 0-40 μM zinc in non-diluted human serum and thus spanning a broad range of clinically relevant concentrations to detect zinc deficiency and toxicity. The common clinical reference range for zinc deficiency is between 8.5-11.5 μM (or 2.1-2.9 μM in 25% serum)^4, 24^. Since our assay can accurately measure zinc in this range, our approach can be easily deployed for a quantitative micronutrient monitoring test at home or in resource-limited environments.

We further highlight that endogenous glucose present in human serum did not interfere with our PGM readout since the metabolic reactions active in cell-free lysate readily removed serum glucose without impacting protein production (**Figure S2A-C**). Previous efforts using PGMs for sensing in serum matrices have required additional steps to remove the confounder of serum glucose from reporter measurements^14, 15^, since serum glucose would be expected to vary from patient to patient—the very reason the PGM exists—and thus interfere with quantitative interpretation of the biosensor readout. Some of the previously reported approaches to solving this problem would be infeasible for practical field application. With our approach, the glucose consumption that was initially a potential obstacle in the use of lysate-based CFE has been repurposed as a distinct advantage that allows quantification across highly variable patient sample matrices.

Furthermore, our strategy of using a transcription factor-based, small molecule inducible genetic circuit is modular and easily generalizable to detect other small molecule targets. To develop a quantitative assay for another target molecule, one simply needs to replace the transcription factor and promoter and adjust the reaction conditions to achieve the desired output level. This only needs to be done once for each new sensor being developed, and is typically best accomplished via systematic variation of plasmid levels to achieve the desired dynamic range of output in the same short assay time, followed by variation of CFE reaction time and/or lactose conversion time if necessary to further increase output signal. The quantification workflow stays unchanged and remains robust to complex samples.

### Extending PGM applications to the detection of bacterial infections

We next tested if PGM-mediated analyte quantification could be extended to quantify targets other than small molecules, as well as if the approach could use genetic circuits based on regulators other than transcription factors. We chose to use toehold switches recognizing RNA sequences of Shiga toxins 1 and 2 (Stx1 and Stx2) to detect pathogenic *E. coli* due to the clinical relevance of the problem^26^ and our previous work in developing these switches^27^. Toehold switches work by RNA-RNA strand hybridization and displacement^28^ for sensitive and fairly specific detection of target sequences. The addition of a trigger sequence complementary to the toehold and partial stem region of the switch unwinds the inhibitory switch hairpin that would otherwise block translation, thereby allowing reporter enzyme expression (**Figure 3A**).

**Figure 3:**
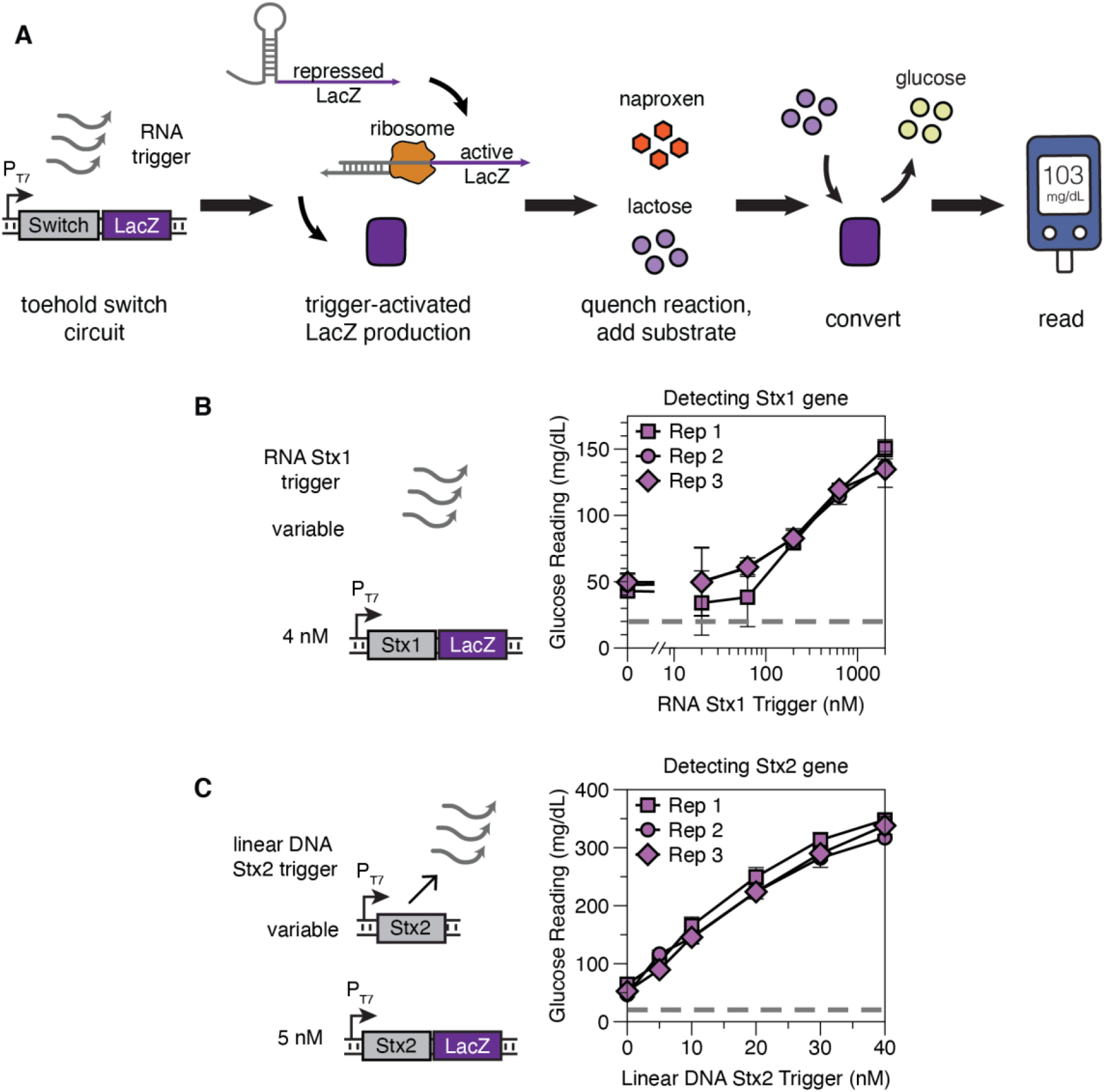
Application of PGM-mediated quantification of nucleic acids in CFE reactions. **(A)** Schematic of toehold switch-modulated LacZ production and LacZ-catalyzed lactose conversion to glucose output. Following 45 min of LacZ production caused by RNA trigger activating a toehold switch to allow translation of LacZ, a mixture of the naproxen-lactose solution was added to quench the CFE reaction and to start lactose conversion for 15 min. The converted glucose was then read on the PGM for target analyte quantification. **(B)** Activation of Stx1 toehold switch and glucose output by RNA Stx1 trigger. Linear glucose response was observed with a logarithmic increment of RNA triggers from 20 to 2000 nM. The same experiment was replicated on different days to verify consistency in glucose output. Replicates (Rep) represent independently assembled reactions and error bars represent the standard deviation of cell-free reaction triplicates in each replicate. Dashed gray line represents PGM’s lowest reading threshold, 20 mg/dL. **(C)** Activation of Stx2 toehold switch and glucose output by linear DNA coding for Stx2 trigger, which can transcribe Stx2 RNA trigger in CFE reaction. Linear glucose response was observed with linear increments of DNA Stx2 trigger from 5 to 40 nM. Replicates (Rep) represent independently assembled reactions and error bars represent the standard deviation of cell-free reaction triplicates in each replicate. Dashed gray line represents PGM’s lowest reading threshold, 20 mg/dL.

Using the same strategy and assay times, we first demonstrated RNA-modulated glucose output over time using an Stx1 toehold switch with a LacZ reporter (**Figure 3B,** see **Supplemental Method** for trigger preparation). DNA of Stx1 triggers was amplified from genomic DNA of Shiga Toxin producing *E. coli* O157:H7 and served as a template for *in vitro* transcription to produce RNA Stx1 triggers. A linear increase in glucose output was observed over logarithmic increments of RNA Stx1 triggers ranging from 20 nM to 2 μM, behavior consistent with RNA trigger-activated toehold switch output in previous reports^3, 4, 29^. Because RNA could also be made from a linear DNA template coding for trigger transcription, we next tested if adding linear DNA could modulate glucose production. We added linear DNA encoding RNA Stx2 trigger amplified from genomic DNA of *E. coli* O157:H7 to activate an Stx2 toehold switch with a LacZ reporter. We observed higher glucose conversion and a lower detection limit than for Stx1 (**Figure 3C**). Compared to the zinc sensor, the toehold switches used here exhibited higher background leakiness, as evidenced by the baseline (0 nM) readouts being approximately 50 mg/dL instead of the PGM’s minimum reading (20 mg/dL). This increased background is due to the use of a dialyzed CFE lysate with enhanced transcriptional activity for this application, as well as due to optimizing switch plasmid concentrations for improved fold-change in glucose readings over the range of trigger concentrations tested, both intentional design decisions for this sensor. It is important to assess the background noise level during the development of any new PGM-based sensor and then compensate for this noise via reaction optimization to whatever extent is feasible. Ideally, background noise would be below the PGM’s detection threshold, and reduction of sensor concentration, reaction time, or lactose substrate concentrations could help achieve this goal, although sometimes other design goals may conflict with this objective and constraint those reaction parameters. Alternatively, a t-test could be done during development to establish the limit of detection for distinguishing signal from noise, allowing identification of the corresponding PGM readings that indicate only background noise. Nevertheless, our results demonstrate that a lysate-based CFE reaction coupled to PGM quantification is a highly generalizable platform compatible with multiple types of analyte inputs and multiple types of genetic regulators.

Although our current nucleic acid sensors could not detect targets at physiologically relevant concentrations (typically attomolar to femtomolar levels), an upstream amplification step can be implemented to bring initial nucleic acid concentrations up to the detection limit^3, 7, 29^. Previous work has shown robust concentration-dependent toehold switch activation with femto- to pico-molar of triggers amplified via isothermal amplification techniques^29^. Further, we note that although for detection of infectious diseases (such as COVID-19, Zika, and Ebola virus), a binary yes/no result may often be sufficient for diagnosis^2, 3, 30^, there are many cases where continuous monitoring and quantification of viral load is essential for assessing treatment efficacy and determining disease prognosis^31^. Having a low-cost, portable, and reliable quantification device can empower patients and healthcare workers to make faster and better medical decisions at the point of need.

## Conclusion

Our work demonstrates that CFE lysate-based biosensors can be easily coupled to a PGM for rapid and reliable analyte quantification at the point of need. The resulting platform is highly generalizable, with demonstrated compatibility with different genetic regulators, analyte types, and sample matrices. Previous efforts repurposing PGMs for small molecule quantification have often used invertase-conjugated antibodies and DNA oligomers, or reagent drop-out methods^8–15^, which have limitations in sensor sensitivity, specificity, and generalizability toward the detection of other targets. Here we show that interfacing synthetic circuits in CFE reactions adds an extensive library of developed biosensors to PGM sensor design and allows users to fine-tune individual sensing reactions with genetic regulators and signaling cascades. Indicative of the modularity and generalizability of this approach, each of the sensing circuits used here was originally developed in the context of different biosensors, but all could be directly integrated into a PGM-based readout with some basic assay optimization (e.g., adjustment of plasmid concentrations and reaction time). The modular utility of the transcription and translation reactions used to transduce signals in CFE reactions means that diverse target analytes can be detected with high sensitivity and specificity while providing critical quantification.

We further highlight that the CFE-based approach is an enabling platform for improved test accessibility and use at the point of need. CFE biosensing reactions have been demonstrated to retain their function after lyophilization^2–4, 27, 32, 33^, including those based on lysates. As a result, these tests can be stored and shipped to testing sites without the cold-chain requirements of PGM-mediated diagnostics that use invertase conjugated antibody or aptamer approaches^10–15^, significantly enabling CFE-based diagnostics’ deployment to the point of need. Moreover, the use of lysate-based CFE in this work rather than purified protein expression systems (like PURExpress) can reduce the cost of CFE reagents by almost an order of magnitude^2, 7^, making such an approach more feasible for wide-scale deployment and accessible to the developing world as well as to consumers in developed countries. We also show that CFE metabolism can be exploited to remove endogenous glucose initially present in complex samples (like human serum) in a one-pot format, thereby eliminating an upstream processing step to remove endogenous glucose that would otherwise be a requirement of a PGM-based method. At the point of use, the operator simply needs to rehydrate the freeze-dried test reaction with sampled fluid to activate the sensing reaction, incubate the sample for a set amount of time, and then add the naproxen-lactose solution to shift the reaction to glucose production. A commercial PGM strip would be used to measure the glucose produced, immediately generating a numerical output on the PGM for result interpretation.

It is worth noting that for successful field deployment of these PGM-based sensors, additional investigation will be necessary. One area that such investigations should focus on is the reduction of batch-to-batch variability in crude lysate and reagent preparation. Interlaboratory variability in cell-free lysate, reagent, and reaction preparation is known to potentially have an impact on quantitative outputs of cell-free reactions^34^. When we assessed the PGM’s ability to accurately quantify targets using cell-free reactions from five crude lysate batches, the inter-lysate variability was found to be low, though the variability due to batches of cell-free reagents was a bit higher (Figure S3). This suggests the importance of standardization, quality assurance, and quality control if this process were to be scaled up for manufacturing and distributed use. Another area worth further investigation is the sensitivity of assay results to perturbations in the time and temperature during the LacZ production and glucose conversion steps. If sensitivity to such perturbations is high, one potential mitigating strategy would be to use a set of standards that could be run in parallel with the test reaction for additional validation^4^; we note that some PGMs already require some degree of calibration by the user. Another possibility could be to engineer a companion device to automate reagent dispensing and regulate reaction temperature and timing. Prototypes for automated, sequential introduction of reagents^35, 36^ and portable incubators^7^ have been developed by different labs for point-of-care diagnostics. Though these additions would increase the capital cost of the equipment and the cost per assay, thus limiting its potential use in low-resource environments, it would still be sufficiently accessible for patient at-home use and clinic use in the developed world.

Nevertheless, our work provides an enabling advance toward inexpensive, point-of-care sample quantification with simpler transportation and operator requirements and fast result turnaround in 1 hr. We demonstrate that interfacing synthetic biology and CFE to PGM-mediated analyte detection has the potential to enable accessible, affordable, and reliable quantification of diverse analytes at the point of need.

## Materials and Methods

### Bacterial Strains and Plasmid Preparation

*E. coli* strain DH10β was used for all cloning and plasmid preparations. *E. coli* strain BL21 Star (DE3) Δ*lacIZYA* was created by lambda red recombination and used for in-house cell-free lysate preparation. Genomic DNA from *E. coli* O157: H7 (ATCC 51657GFP) was used as a template for Stx1 and Stx2 trigger amplification.

Supplementary **Table S1** contains sequences of all parts used in this study. Eurofins Genomics synthesized DNA oligonucleotides for cloning and sequencing. Plasmid DNA used for all CFE reactions was purified from EZNA midiprep columns (OMEGA Bio-Tek) followed by isopropanol and ethanol precipitation. The purified DNA pellets were reconstituted in the elution buffer, measured on a Nanodrop 2000 for concentration, and stored at −20 °C until use.

### Cell-Free Reactions

The cell-free reaction composition was as previously described by Kwon and Jewett^37^. All lysate and reagent master mixes were thawed on ice and had less than 3 freeze-thaw cycles. All reactions were assembled on ice and in PCR tube strips. The reaction master mix containing lysate was vortexed at a medium-high setting to ensure homogenous mixing before being aliquoted into individual reactions. See **Supplemental Method** for details on the crude cell-free extract preparation. All CFE reactions, except reactions expressing toehold switches or malachite green aptamers, used crude lysate without post lysate processing steps such as run-off reactions and dialysis. Dialyzed lysate was used for toehold switch and malachite green aptamer reactions due to its enhanced transcriptional capacity^38^. **Table S2** tabulates the specified concentrations of plasmids and reaction additives used in each figure.

For reactions measured with a plate reader, each cell-free reaction was 10 μL in volume and pipetted into a black-bottomed 384-well plate (Greiner Bio-One) for fluorescence measurement or a clear-bottomed 384-well plate (Greiner Bio-One) for absorbance measurement. Kinetic reads were performed in a plate reader (Synergy4, BioTek) at 37 °C for 1 hr. The filter setting for GFP measurement was 485/510 nm excitation/emission wavelengths, with the gain set at 70. The filter setting for malachite green measurement was 615/650 nm excitation/emission wavelengths, with the gain set at 100. For chlorophenol red-β-D-galactopyranoside (CPRG) measurement, sample absorbance was measured at 580 nm. All plates were sealed with a transparent, adhesive film to prevent evaporation.

For reactions read on a PGM, each assembled cell-free reaction was 9 μL in volume and placed in a PCR tube with the cap on to prevent evaporation. Reactions were incubated at 37 °C in a thermocycler for the specified amount of time before glucose measurement. For reactions quenched with the naproxen-lactose mix, 1 μL of the 10x quench mix (100 mM naproxen sodium and 400 mM lactose) was added to each reaction and the mixture was vortexed at medium-high setting, settled to the bottom of the tube using a mini centrifuge, and incubated at 37 °C for 15 minutes before measurement on a PGM. The same concentration of the naproxen-lactose mix was added to all cell-free reactions.

### PGM Quantification

A glucose oxidase-based PGM (OneTouch Ultra 2 Blood Glucose Monitoring System, LifeScan Inc) and accompanying test strips (OneTouch Ultra Test Strips, LifeScan Inc) were used for glucose measurement. Once the glucose-generating step of the reaction was completed, 2 μL of each reaction was spotted on the test strip and measured with the PGM. Because the PGM’s readout range is from 20 to 600 mg/dL, values below or above the meter threshold were assigned a value of 20 mg/dL or 600 mg/dL, respectively.

## Acknowledgment

We thank Dr. Michael Jewett for his gift of pJL1 (P_T7_-sfGFP) plasmid. We thank Dr. Julius Lucks for his gift of the plasmid encoding malachite green aptamer (P_T7_-MGA). We thank Dr. Shuichi Takayama for his gift of genomic DNA of *E. coli* O157: H7 (ATCC 51657GFP).

## Author Contributions

Conceptualization: YZ, MPS; Investigation: YZ, PLS, MWK; Formal Analysis: YZ, PLS, MWK; Writing – Original Draft: YZ; Writing – Review & Editing: YZ, PLS, MWK, MPS; Visualization: YZ; Supervision: MPS; Funding Acquisition: MPS.

## Supporting Information

The included supporting information file contains methods for cell-free lysate, human serum, and toehold switch trigger preparation; supplementary figures S1-S4 showing characterization of naproxen quenching in CFE reactions, endogenous metabolism in CFE reactions readily removing a wide range of glucose spiked into human serum without impacting protein expression, assessment of lysate batch-to-batch differences in target quantification, and specificity assessment of primers designed to amplify either Stx1 or Stx2 triggers from genomic DNA; and supplementary tables S1-S3 describing plasmid parts and DNA sequences in this paper, the contents present in CFE reactions in each figure, and the primers used for trigger DNA amplification from *E. coli* O157: H7 genomic DNA template.

## Reference

1. Silverman, A. D.; Karim, A. S.; Jewett, M. C., Cell-free gene expression: an expanded repertoire of applications. Nat Rev Genet 2020, 21 (3), 151–170.

2. Pardee, K.; Green, A. A.; Ferrante, T.; Cameron, D. E.; DaleyKeyser, A.; Yin, P.; Collins, J. J., Paper-based synthetic gene networks. Cell 2014, 159 (4), 940–54.

3. Pardee, K.; Green, A. A.; Takahashi, M. K.; Braff, D.; Lambert, G.; Lee, J. W.; Ferrante, T.; Ma, D.; Donghia, N.; Fan, M.; Daringer, N. M.; Bosch, I.; Dudley, D. M.; O’Connor, D. H.; Gehrke, L.; Collins, J. J., Rapid, Low-Cost Detection of Zika Virus Using Programmable Biomolecular Components. Cell 2016, 165 (5), 1255–1266.

4. McNerney, M. P.; Zhang, Y.; Steppe, P.; Silverman, A. D.; Jewett, M. C.; Styczynski, M. P., Point-of-care biomarker quantification enabled by sample-specific calibration. Sci Adv 2019, 5 (9), eaax4473.

5. Jung, J. K.; Alam, K. K.; Verosloff, M. S.; Capdevila, D. A.; Desmau, M.; Clauer, P. R.; Lee, J. W.; Nguyen, P. Q.; Pasten, P. A.; Matiasek, S. J.; Gaillard, J. F.; Giedroc, D. P.; Collins, J. J.; Lucks, J. B., Cell-free biosensors for rapid detection of water contaminants. Nat Biotechnol 2020.

6. Clarke, S. F.; Foster, J. R., A history of blood glucose meters and their role in self-monitoring of diabetes mellitus. Br J Biomed Sci 2012, 69 (2), 83–93.

7. Amalfitano, E.; Karlikow, M.; Norouzi, M.; Jaenes, K.; Cicek, S.; Masum, F.; Sadat Mousavi, P.; Guo, Y.; Tang, L.; Sydor, A.; Ma, D.; Pearson, J. D.; Trcka, D.; Pinette, M.; Ambagala, A.; Babiuk, S.; Pickering, B.; Wrana, J.; Bremner, R.; Mazzulli, T.; Sinton, D.; Brumell, J. H.; Green, A. A.; Pardee, K., A glucose meter interface for point-of-care gene circuit-based diagnostics. Nat Commun 2021, 12 (1), 724.

8. Jang, Y. J.; Lee, K. H.; Yoo, T. H.; Kim, D. M., Interfacing a Personal Glucose Meter with Cell-Free Protein Synthesis for Rapid Analysis of Amino Acids. Anal Chem 2019, 91 (3), 2531–2535.

9. Lisi, F.; Peterson, J. R.; Gooding, J. J., The application of personal glucose meters as universal point-of-care diagnostic tools. Biosens Bioelectron 2020, 148, 111835.

10. Xiang, Y.; Lan, T.; Lu, Y., Using the widely available blood glucose meter to monitor insulin and HbA1c. J Diabetes Sci Technol 2014, 8 (4), 855–8.

11. Xiang, Y.; Lu, Y., Using personal glucose meters and functional DNA sensors to quantify a variety of analytical targets. Nat Chem 2011, 3 (9), 697–703.

12. Xiang, Y.; Lu, Y., Portable and quantitative detection of protein biomarkers and small molecular toxins using antibodies and ubiquitous personal glucose meters. Anal Chem 2012, 84 (9), 4174–8.

13. Xiang, Y.; Lu, Y., Using commercially available personal glucose meters for portable quantification of DNA. Anal Chem 2012, 84 (4), 1975–80.

14. Yan, L.; Zhu, Z.; Zou, Y.; Huang, Y.; Liu, D.; Jia, S.; Xu, D.; Wu, M.; Zhou, Y.; Zhou, S.; Yang, C. J., Target-responsive “sweet” hydrogel with glucometer readout for portable and quantitative detection of non-glucose targets. J Am Chem Soc 2013, 135 (10), 3748–51.

15. Zhang, J.; Xiang, Y.; Wang, M.; Basu, A.; Lu, Y., Dose-Dependent Response of Personal Glucose Meters to Nicotinamide Coenzymes: Applications to Point-of-Care Diagnostics of Many Non-Glucose Targets in a Single Step. Angew Chem Int Ed Engl 2016, 55 (2), 732–6.

16. Combs, G. F., Jr.; Trumbo, P. R.; McKinley, M. C.; Milner, J.; Studenski, S.; Kimura, T.; Watkins, S. M.; Raiten, D. J., Biomarkers in nutrition: new frontiers in research and application. Ann N Y Acad Sci 2013, 1278, 1–10.

17. Goossens, N.; Nakagawa, S.; Sun, X.; Hoshida, Y., Cancer biomarker discovery and validation. Transl Cancer Res 2015, 4 (3), 256–269.

18. Vasan, R. S., Biomarkers of cardiovascular disease: molecular basis and practical considerations. Circulation 2006, 113 (19), 2335–62.

19. Calhoun, K. A.; Swartz, J. R., Energizing cell-free protein synthesis with glucose metabolism. Biotechnol Bioeng 2005, 90 (5), 606–13.

20. Caschera, F.; Noireaux, V., A cost-effective polyphosphate-based metabolism fuels an all E. coli cell-free expression system. Metab Eng 2015, 27, 29–37.

21. Karim, A. S.; Rasor, B. J.; Jewett, M. C., Enhancing control of cell-free metabolism through pH modulation. Synthetic Biology 2019, 5 (1).

22. Miguez, A. M.; McNerney, M. P.; Styczynski, M. P., Metabolic Profiling of Escherichia coli-based Cell-Free Expression Systems for Process Optimization. Ind Eng Chem Res 2019, 58 (50), 22472–22482.

23. Miguez, A. M.; Zhang, Y.; Piorino, F.; Styczynski, M. P., Metabolic Dynamics in *Escherichia coli*-based Cell-Free Systems. bioRxiv 2021, 2021.05.16.444339.

24. King, J. C.; Brown, K. H.; Gibson, R. S.; Krebs, N. F.; Lowe, N. M.; Siekmann, J. H.; Raiten, D. J., Biomarkers of Nutrition for Development (BOND)-Zinc Review. J Nutr 2015, 146 (4), 858S–885S.

25. Bhutta, Z. A.; Das, J. K.; Rizvi, A.; Gaffey, M. F.; Walker, N.; Horton, S.; Webb, P.; Lartey, A.; Black, R. E.; Lancet Nutrition Interventions Review Group, t. M.; Child Nutrition Study, G., Evidence-based interventions for improvement of maternal and child nutrition: what can be done and at what cost? Lancet 2013, 382 (9890), 452–477.

26. Karmali, M. A., Infection by Shiga toxin-producing Escherichia coli: an overview. Mol Biotechnol 2004, 26 (2), 117–22.

27. Zhang, Y.; Kojima, T.; Kim, G.-A.; McNerney, M. P.; Takayama, S.; Styczynski, M. P., Protocell Arrays for Simultaneous Detection of Diverse Analytes. bioRxiv 2021, 2021.02.13.431022.

28. Green, A. A.; Silver, P. A.; Collins, J. J.; Yin, P., Toehold switches: de-novo-designed regulators of gene expression. Cell 2014, 159 (4), 925–39.

29. Takahashi, M. K.; Tan, X.; Dy, A. J.; Braff, D.; Akana, R. T.; Furuta, Y.; Donghia, N.; Ananthakrishnan, A.; Collins, J. J., A low-cost paper-based synthetic biology platform for analyzing gut microbiota and host biomarkers. Nat Commun 2018, 9 (1), 3347.

30. Carter, L. J.; Garner, L. V.; Smoot, J. W.; Li, Y.; Zhou, Q.; Saveson, C. J.; Sasso, J. M.; Gregg, A. C.; Soares, D. J.; Beskid, T. R.; Jervey, S. R.; Liu, C., Assay Techniques and Test Development for COVID-19 Diagnosis. ACS Cent Sci 2020, 6 (5), 591–605.

31. Berger, A.; Braner, J.; Doerr, H. W.; Weber, B., Quantification of viral load: clinical relevance for human immunodeficiency virus, hepatitis B virus and hepatitis C virus infection. Intervirology 1998, 41 (1), 24–34.

32. Liu, X.; Silverman, A. D.; Alam, K. K.; Iverson, E.; Lucks, J. B.; Jewett, M. C.; Raman, S., Design of a Transcriptional Biosensor for the Portable, On-Demand Detection of Cyanuric Acid. ACS Synth Biol 2020, 9 (1), 84–94.

33. Thavarajah, W.; Silverman, A. D.; Verosloff, M. S.; Kelley-Loughnane, N.; Jewett, M. C.; Lucks, J. B., Point-of-Use Detection of Environmental Fluoride via a Cell-Free Riboswitch-Based Biosensor. ACS Synth Biol 2020, 9 (1), 10–18.

34. Cole, S. D.; Beabout, K.; Turner, K. B.; Smith, Z. K.; Funk, V. L.; Harbaugh, S. V.; Liem, A. T.; Roth, P. A.; Geier, B. A.; Emanuel, P. A.; Walper, S. A.; Chavez, J. L.; Lux, M. W., Quantification of Interlaboratory Cell-Free Protein Synthesis Variability. ACS Synth Biol 2019, 8 (9), 2080–2091.

35. Gayet, R. V.; de Puig, H.; English, M. A.; Soenksen, L. R.; Nguyen, P. Q.; Mao, A. S.; Angenent-Mari, N. M.; Collins, J. J., Creating CRISPR-responsive smart materials for diagnostics and programmable cargo release. Nat Protoc 2020, 15 (9), 3030–3063.

36. English, M. A.; Soenksen, L. R.; Gayet, R. V.; de Puig, H.; Angenent-Mari, N. M.; Mao, A. S.; Nguyen, P. Q.; Collins, J. J., Programmable CRISPR-responsive smart materials. Science 2019, 365 (6455), 780–785.

37. Kwon, Y. C.; Jewett, M. C., High-throughput preparation methods of crude extract for robust cell-free protein synthesis. Sci Rep-Uk 2015, 5.

38. Silverman, A. D.; Kelley-Loughnane, N.; Lucks, J. B.; Jewett, M. C., Deconstructing Cell-Free Extract Preparation for in Vitro Activation of Transcriptional Genetic Circuitry. ACS Synth Biol 2019, 8 (2), 403–414.

