## Supplementary figures and information for "Point-of-care analyte quantification and digital readout via lysate-based cell-free biosensors interfaced with personal glucose monitors"

School of Chemical and Biomolecular Engineering, Georgia Institute of Technology, Atlanta, GA, 30332-0100, United States.
* These authors contributed equally to this work
† Corresponding author

**Table of Contents**

Supplemental Method: ............ S2

*Crude E. coli Cell-Free Lysate Preparation* ............ S2

*Serum Processing* ............ S2

*Trigger Preparation* ............ S2

Figure S1: Characterization of naproxen quenching in CFE reactions. ............ S4

Figure S2: Endogenous metabolism in CFE reactions readily removed a wide range of glucose spiked into human serum without impacting protein expression. ............ S6

Figure S3: Assessment of lysate batch-to-batch variability in target quantification. ............ S7

Figure S4: Specificity assessment of primers designed to amplify either *stx1* or *stx2* triggers from genomic DNA. ............ S8

Table S1: Description of plasmid parts and DNA sequences in this paper. ............ S9

Table S2. Description of lysate, plasmid concentrations, and reaction additives present in CFE reactions in each figure. ............ S13

Table S3: Primers used for trigger DNA amplification from *E. coli* O157: H7 genomic DNA template. ............ S14

References: ............ S16

**Supplemental Method:**

*Crude E. coli Cell-Free Lysate Preparation*

Cellular lysate for all experiments was prepared as described by Sun *et al.*[1] with a few protocol modifications. Briefly, BL21 Star (DE3) Δ*lacIZYA* cells were grown in 2× YTP medium at 37 ˚C and 220 rpm to an optical density (OD) of 0.3-0.5 for IPTG induction at 0.4 mM to activate expression of T7 RNAP Polymerase. The cells were grown further until the OD was 1.5-2.0, corresponding to the mid-exponential growth phase. Cells were centrifuged at 2700 rcf and washed three times with S30A buffer (50 mM tris, 14 mM magnesium glutamate, 60 mM potassium glutamate, 2 mM dithiothreitol, and pH-corrected to 7.7 with acetic acid). After the final centrifugation, the wet cell mass was determined, and cells were resuspended in 1 mL of S30A buffer per 1 g of wet cell mass. The cellular resuspension was divided into 1 mL aliquots. Cells were lysed using a Q125 sonicator (Qsonica) at a frequency of 20 kHz and 50% amplitude. Cells were sonicated on ice with cycles of 10 s on and 10 s off, delivering approximately 200 J, at which point the cells appeared visibly lysed. An additional 4 mM dithiothreitol was added to each tube, and the sonicated mixture was then centrifuged at 12,000 rcf and 4 ˚C for 10 min. After centrifugation, the supernatant was removed, divided into 100 $\mu$L aliquots, and stored at -80 ˚C until use.

An additional runoff reaction and dialysis were performed in lysate used for expression of toehold switches and malachite green aptamer. Briefly, the centrifuged sonication product was incubated at 37 ˚C and 220 rpm for 80 min. After this runoff reaction, the cellular lysate was centrifuged at 12,000 rcf and 4 ˚C for 10 min. The supernatant was removed and loaded into a 10 kDa molecular weight cutoff dialysis cassette (Thermo Fisher). The lysate was dialyzed in 1 liter of S30B buffer (14 mM magnesium glutamate, 60 mM potassium glutamate, 1 mM dithiothreitol, and pH-corrected to 8.2 with tris) at 4 ˚C for 3 hours. Dialyzed lysate was removed and centrifuged at 12,000 rcf and 4 ˚C for 10 min. The supernatant was removed, aliquoted in volumes of 100 $\mu$L, and stored at -80 ˚C for future use.

*Serum Processing*

Pooled human serum was collected from donors as approved in Institutional Review Board protocol number H17489. Venous blood was collected in 6 mL of BD Vacutainer collection tubes for trace element testing, and tubes were left on ice for about 30 min to clot. Blood was transferred to a 50 mL conical tube and centrifuged at 2700 rcf for 30 min at 4 ˚C. Serum was removed, and an aliquot of untreated human serum was saved for zinc baseline analysis. The remaining serum was treated with Chelex 100 resin by adding 1 mg resin per 1 mL of serum and vigorously stirred for 2 hrs at room temperature. The resin was isolated from samples through centrifugation and syringe filtering. All serum samples were aliquoted to minimize free-thaw cycles and stored at -20 ˚C until use.

Measurement of successful zinc removal from serum was done at the University of Georgia Laboratory for Environmental Analysis (**Figure S2D**). Samples were digested with concentrated acid and analyzed on ICP-MS according to EPA method 3052.

*Trigger Preparation*

DNA encoding each trigger RNA used in experiments was amplified from the genomic DNA of *E. coli* O157: H7 *via* PCR with Q5 DNA polymerase (New England Biolabs). Sequences for primers used to amplify triggers from DNA template or genomic DNA are provided in **Table S3**. After PCR amplification, products were run on a 2 w/v% agarose gel to verify strain-specific amplification of targets (**Figure S4**) and then purified using a PCR purification kit (Omega Bio-

Tek). The prepared linear DNA was either directly used in cell-free reactions or used as a template for *in vitro* transcription.

RNA triggers were transcribed from a linear DNA template using T7 polymerase according to the manufacturer's protocol (New England Biolabs). Following RNA synthesis, Dnase I (Zymo Research) was added to degrade the linear DNA template. The RNA products were then purified using an RNA Clean and Concentrator kit (Zymo Research) according to the manufacturer's protocol. Following purification, RNA concentration was measured on a Nanodrop 2000, aliquoted, and stored at -20 ˚C.

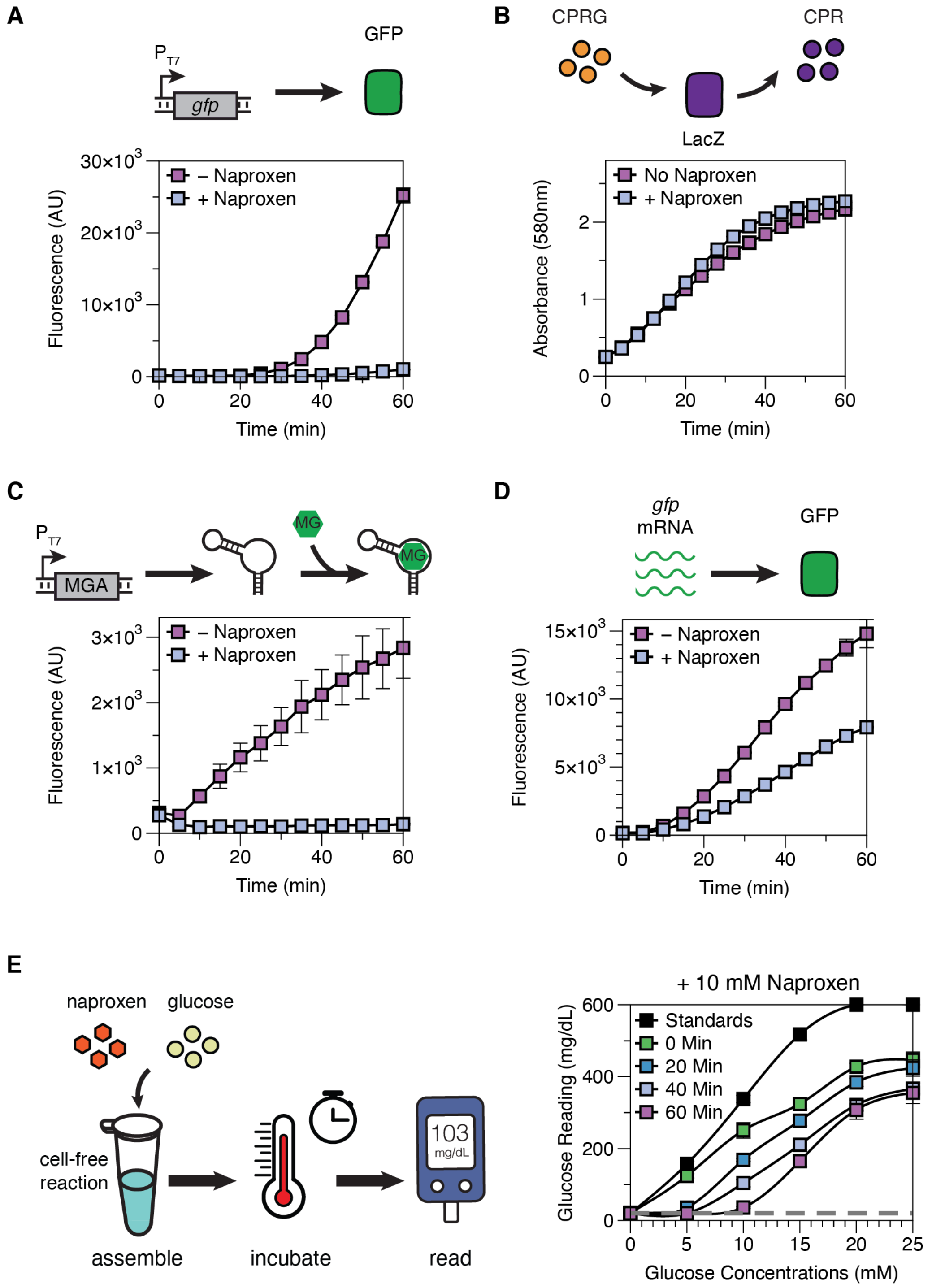


**Figure S1:** Characterization of naproxen quenching in CFE reactions. Error bars in each sub-panel represent standard deviations of cell-free reaction triplicates. **(A)** Plasmid encoding for GFP expression was added to each reaction for fluorescent signal production to characterize naproxen's cumulative impact on protein transcription and translation. Addition of naproxen sodium at the beginning of the CFE reaction inhibits GFP expression. **(B)** BL21 Star (DE3)

lysate containing LacZ enzyme was dosed into a CFE reaction at 5% volume. LacZ activity was measured by its ability to cleave chlorophenol red-β-D-galactopyranoside (CPRG) to chlorophenol red. Addition of naproxen sodium did not significantly impact LacZ activity. **(C)** Plasmid encoding for malachite green RNA aptamer (MGA) expression and malachite green dye (MG) were added to each reaction for fluorescent signal production to characterize transcription. Addition of naproxen sodium at the beginning of the CFE reaction prevents transcription. **(D)** RNA transcripts coding for GFP translation were added to each CFE reaction for fluorescent signal production. Addition of naproxen sodium resulted in ~46% repression of translation. **(E)** Addition of 10 mM naproxen slowed down endogenous glucose consumption in CFE reactions, with glucose readings (especially at later time points) higher in the presence of naproxen than in **Figure 1A**. Standards presented here are the same set of data presented in **Figure 1A**.

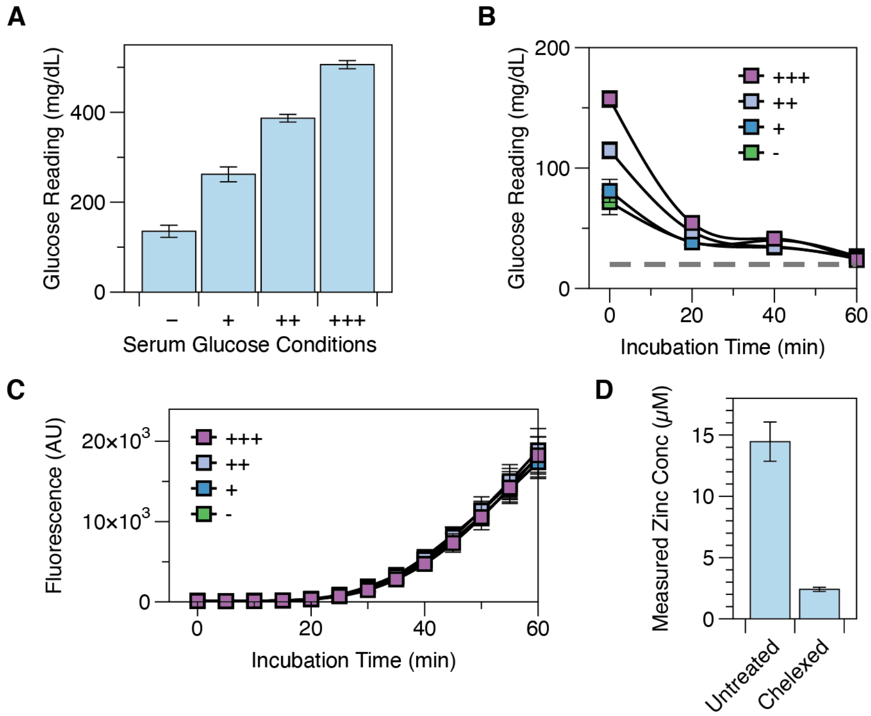


**Figure S2:** Endogenous metabolism in CFE reactions readily removed a wide range of glucose spiked into human serum without impacting protein expression. **(A)** Glucose was added to human serum to span the full detection range of the glucose monitor (OneTouch Ultra 2). Error bars in each sub-panel represent standard deviations of technical triplicates. **(B)** Human serum containing various amounts of glucose was added to a CFE reaction (with no LacZ expression) at 25% volume, and significant depletion of glucose was observed in the first 20 minutes for all reactions. Error bars in each sub-panel represent standard deviations of reaction triplicates. **(C)** GFP expression was unaffected in CFE reactions containing 25% human serum at a wide range of glucose concentrations. Error bars in each sub-panel represent standard deviations of two independently assembled experimental replicates, each with cell-free reaction triplicates. **(D)** Verification via ICP-MS of serum zinc removal after treatment with Chelex-100 resin.

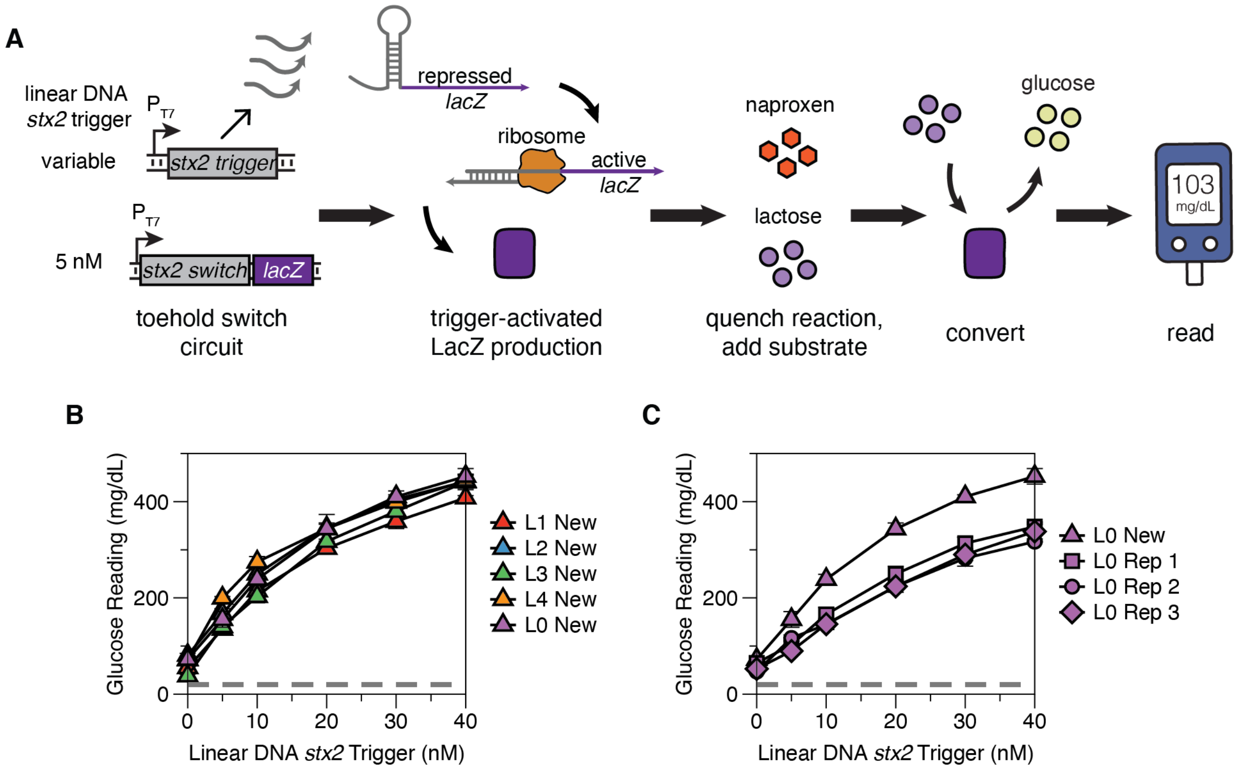


**Figure S3:** Assessment of lysate batch-to-batch variability in target quantification. **(A)** Schematic of toehold switch sensor used and process workflow. Linear DNA expressing the *stx2* trigger sequence was added at 0-40 nM to a cell-free reaction containing 5 nM of the cognate *stx2* toehold switch. Experiment workflow is the same as **Figure 3C**: reactions were incubated at 37 ˚C for 45 min before being quenched by the naproxen-lactose mix and incubated for another 15 min before PGM measurement. **(B)** Comparison of linear DNA quantification using different batches of lysate prepared using the same protocol. Low inter-lysate variability was observed across five batches of lysates. L0-2 were prepared over the course of 2019, and L3-4 were prepared in spring 2021. The “New” designation in the figure legend indicates a different batch of cell-free reagent was used in data collection compared to that used for the remainder of the figures (L0). Error bars represent the standard deviation of cell-free reaction triplicates. Dashed gray line represents PGM’s lowest reading threshold, 20 mg/dL. **(C)** Comparison of linear DNA quantification in the same lysate batch but different reagent batches or aliquots. L0 Rep 1-3 are the same set of data shown in **Figure 3C** using old reagents, while L0 New is the same lysate using a new batch of reagents. While the trend of signal production is preserved, the new batch of reagent generated higher glucose readings for the same amount of linear DNA input. Thus, there is the potential for batch-to-batch variability at multiple levels in these reactions that should be controlled for carefully.

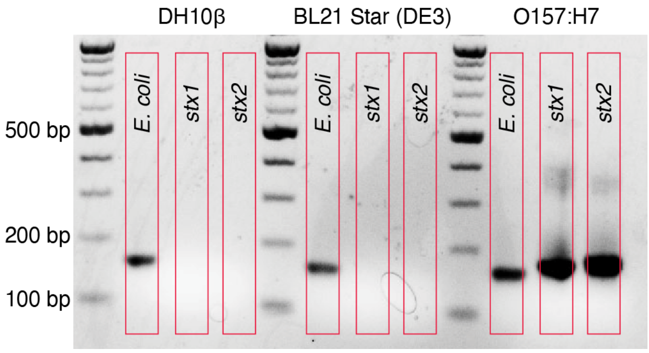


**Figure S4:** Specificity assessment of primers designed to amplify either *stx1* or *stx2* triggers from genomic DNA templates. *stx1* and *stx2* trigger DNA were strongly amplified when the appropriate template (*E. coli* O157:H7 genomic DNA from ATCC 51657GFP) was added, and no amplification of *stx1* and *stx2* triggers was observed from the genomic DNA of common lab *E. coli* strains DH10β or BL21 Star (DE3). A previously developed primer pair recognizing common *E. coli* strains[2] was added as a positive control.

**Table S1:** Description of plasmid parts and DNA sequences in this paper.

<table>
<tr><th>Name</th><th>Construct Description</th></tr>
<tr><td>$P_{T7}$-lacZ</td><td>Plasmid encoding β-galactosidase (LacZ) expression under the T7 promoter</td></tr>
<tr><td colspan="2">T7 Promoter- Stability Hairpin-StrongRBS-lacZ-T7 Terminator<br><br>taatacgactcactatagggagaccacaacggtttccctctagaaataattttgtttaactttaagaaggagatatacatatgACCATGATTACGGATTCACTGGCCGT<br>CGTTTTACAACGTCGTGACTGGGAAAACCCTGGCGTTACCCAACTTAATCGCCTTGCAGCACATCCCCCTTTCGCCAGCTGG<br>CGTAATAGCGAAGAGGCCCGCACCGATCGCCCTTCCCAACAGTTGCGCAGCCTGAATGGCGAATGGCGCTTTGCCTGGTTT<br>CCGGCACCAGAAGCGGTGCCGGAAAGCTGGCTGGAGTGCGATCTTCCTGAGGCCGATACTGTCGTCGTCCCCTCAAACTG<br>GCAGATGCACGGTTACGATGCGCCCATCTACACCAACGTGACCTATCCCATTACGGTCAATCCGCCGTTTGTTCCCACGGA<br>GAATCCGACGGGTTGTTACTCGCTCACATTTAATGTTGATGAAAGCTGGCTACAGGAAGGCCAGACGCGAATTATTTTTGAT<br>GGCGTTAACTCGGCGTTTCATCTGTGGTGCAACGGGCGCTGGGTCGGTTACGGCCAGGACAGTCGTTTGCCGTCTGAATTT<br>GACCTGAGCGCATTTTTACGCGCCGGAGAAAACCGCCTCGCGGTGATGGTGCTGCGCTGGAGTGACGGCAGTTATCTGGA<br>AGATCAGGATATGTGGCGGATGAGCGGCATTTTCCGTGACGTCTCGTTGCTGCATAAACCGACTACACAAATCAGCGATTTC<br>CATGTTGCCACTCGCTTTAATGATGATTTCAGCCGCGCTGTACTGGAGGCTGAAGTTCAGATGTGCGGCGAGTTGCGTGACT<br>ACCTACGGGTAACAGTTTCTTTATGGCAGGGTGAAACGCAGGTCGCCAGCGGCACCGCGCCTTTCGGCGGTGAAATTATCG<br>ATGAGCGTGGTGGTTATGCCGATCGCGTCACACTACGTCTGAACGTCGAAAACCCGAAACTGTGGAGCGCCGAAATCCCGA<br>ATCTCTATCGTGCGGTGGTTGAACTGCACACCGCCGACGGCACGCTGATTGAAGCAGAAGCCTGCGATGTCGGTTTCCGCG<br>AGGTGCGGATTGAAAATGGTCTGCTGCTGCTGAACGGCAAGCCGTTGCTGATTCGAGGCGTTAACCGTCACGAGCATCATC<br>CTCTGCATGGTCAGGTCATGGATGAGCAGACGATGGTGCAGGATATCCTGCTGATGAAGCAGAACAACTTTAACGCCGTGC<br>GCTGTTCGCATTATCCGAACCATCCGCTGTGGTACACGCTGTGCGACCGCTACGGCCTGTATGTGGTGGATGAAGCCAATA<br>TTGAAACCCACGGCATGGTGCCAATGAATCGTCTGACCGATGATCCGCGCTGGCTACCGGCGATGAGCGAACGCGTAACG<br>CGAATGGTGCAGCGCGATCGTAATCACCCGAGTGTGATCATCTGGTCGCTGGGGAATGAATCAGGCCACGGCGCTAATCAC<br>GACGCGCTGTATCGCTGGATCAAATCTGTCGATCCTTCCCGCCCGGTGCAGTATGAAGGCGGCGGAGCCGACACCACGGC<br>CACCGATATTATTTGCCCGATGTACGCGCGCGTGGATGAAGACCAGCCCTTCCCGGCTGTGCCGAAATGGTCCATCAAAAA<br>ATGGCTTTCGCTACCTGGAGAGACGCGCCCGCTGATCCTTTGCGAATACGCCCACGCGATGGGTAACAGTCTTGGCGGTTT<br>CGCTAAATACTGGCAGGCGTTTCGTCAGTATCCCCGTTTACAGGGCGGCTTCGTCTGGGACTGGGTGGATCAGTCGCTGAT<br>TAAATATGATGAAAACGGCAACCCGTGGTCGGCTTACGGCGGTGATTTTGGCGATACGCCGAACGATCGCCAGTTCTGTAT<br>GAACGGTCTGGTCTTTGCCGACCGCACGCCGCATCCAGCGCTGACGGAAGCAAAACACCAGCAGCAGTTTTTCCAGTTCCG<br>TTTATCCGGGCAAACCATCGAAGTGACCAGCGAATACCTGTTCCGTCATAGCGATAACGAGCTCCTGCACTGGATGGTGGC<br>GCTGGATGGTAAGCCGCTGGCAAGCGGTGAAGTGCCTCTGGATGTCGCTCCACAAGGTAAACAGTTGATTGAACTGCCTGA<br>ACTACCGCAGCCGGAGAGCGCCGGGCAACTCTGGCTCACAGTACGCGTAGTGCAACCGAACGCGACCGCATGGTCAGAAG<br>CCGGACACATCAGCGCCTGGCAGCAGTGGCGTCTGGCTGAAAACCTCAGCGTGACACTCCCCGCCGCGTCCCACGCCATC<br>CCGCATCTGACCACCAGCGAAATGGATTTTTGCATCGAGCTGGGTAATAAGCGTTGGCAATTTAACCGCCAGTCAGGCTTTC<br>TTTCACAGATGTGGATTGGCGATAAAAAACAACTGCTGACGCCGCTGCGCGATCAGTTCACCCGTGCACCGCTGGATAACG<br>ACATTGGCGTAAGTGAAGCGACCCGCATTGACCCTAACGCCTGGGTCGAACGCTGGAAGGCGGCGGGCCATTACCAGGCC<br>GAAGCAGCGTTGTTGCAGTGCACGGCAGATACACTTGCTGATGCGGTGCTGATTACGACCGCTCACGCGTGGCAGCATCAG<br>GGGAAAACCTTATTTATCAGCCGGAAAACCTACCGGATTGATGGTAGTGGTCAAATGGCGATTACCGTTGATGTTGAAGTGG<br>CGAGCGATACACCGCATCCGGCGCGGATTGGCCTGAACTGCCAGCTGGCGCAGGTAGCAGAGCGGGTAAACTGGCTCGG<br>ATTAGGGCCGCAAGAAAACTATCCCGACCGCCTTACTGCCGCCTGTTTTGACCGCTGGGATCTGCCATTGTCAGACATGTAT<br>ACCCCGTACGTCTTCCCGAGCGAAAACGGTCTGCGCTGCGGGACGCGCGAATTGAATTATGGCCCACACCAGTGGCGCGG<br>CGACTTCCAGTTCAACATCAGCCGCTACAGTCAACAGCAACTGATGGAAACCAGCCATCGCCATCTGCTGCACGCGGAAGA<br>AGGCACATGGCTGAATATCGACGGTTTCCATATGGGGATTGGTGGCGACGACTCCTGGAGCCCGTCAGTATCGGCGGAATT<br>CCAGCTGAGCGCCGGTCGCTACCATTACCAGTTGGTCTGGTGTCAAAAAtaagtcgaccggctgctaacaaagcccgaaaggaagctgagttgg<br>ctgctgccaccgctgagcaataactagcataaccccttggggcctctaaacgggtcttgaggggttttttg</td></tr>
</table>

<table>
<tr><td>$P_{T7}$-zntR</td><td>Plasmid encoding ZntR expression under T7 promoter with a strong ribosomal binding site</td></tr>
<tr><td colspan="2">T7 Promoter-Stability Hairpin-StrongRBS-zntR-T7 Terminator<br><br>taatacgactcactatagggagaccacaacggtttccctctagaaataattttgtttaactttaagaaggagatatacatATGTATCGCATTGGTGAGCTGGCAAAAT<br>GGCGGAAGTAACACCCGACACGATTCGTTATTACGAAAAACAGCAGATGATGGAGCATGAAGTGCGTACTGAAGGTGGGTT<br>TCGCCTATATACCGAAAGCGATCTCCAGCGATTGAAATTTATCCGCCATGCCAGACAACTAGGTTTCAGTCTGGAGTCGATC<br>CGCGAGTTGCTGTCGATCCGCATCGATCCTGAACACCATACCTGTCAGGAGTCAAAAGGCATTGTGCAGGAAAGATTGCAG<br>GAAGTCGAAGCACGGATAGCCGAGTTGCAGAGTATGCAGCGTTCCTTGCAACGCCTTAACGATGCCTGTTGTGGACTGCT<br>CATAGCAGTGTTTATTGTTCGATTCTTGAAGCTCTTGAACAAGGGGCGAGTGGCGTTAAGAGTGGTTGTTGATAAgtcgaccggct<br>gctaacaaagcccgaaaggaagctgagttggctgctgccaccgctgagcaataactagcataaccccttggggcctctaaacgggtcttgaggggttttttg</td></tr>
</table>

| $P_{ZntA}$-*lacZ* | Plasmid encoding LacZ expression under ZntR regulated $P_{ZntA}$ promoter |
|---|---|

$P_{ZntA}$-Stability Hairpin-StrongRBS- *lacZ*- T7 Terminator

CTGTATCTCTGATAAAACTTGACTCTGGAGTCGACTCCAGAGTGTATCCTTCGGTTAAT
ccacaacggtttccctctagaaataattttgtttaactttaagaaggagatatacatatgACCATGATTACGGATTCACTGGCCGTCGTTTTACAACGTCGTG
ACTGGGAAAACCCTGGCGTTACCCAACTTAATCGCCTTGCAGCACATCCCCCTTTCGCCAGCTGGCGTAATAGCGAAGAGG
CCCGCACCGATCGCCCTTCCCAACAGTTGCGCAGCCTGAATGGCGAATGGCGCTTTGCCTGGTTTCCGGCACCAGAAGCG
GTGCCGGAAAGCTGGCTGGAGTGCGATCTTCCTGAGGCCGATACTGTCGTCGTCCCCTCAAACTGGCAGATGCACGGTTAC
GATGCGCCCATCTACACCAACGTGACCTATCCCATTACGGTCAATCCGCCGTTTGTTCCCACGGAGAATCCGACGGGTTGTT
ACTCGCTCACATTTAATGTTGATGAAAGCTGGCTACAGGAAGGCCAGACGCGAATTATTTTTGATGGCGTTAACTCGGCGTT
TCATCTGTGGTGCAACGGGCGCTGGGTCGGTTACGGCCAGGACAGTCGTTTGCCGTCTGAATTTGACCTGAGCGCATTTTT
ACGCGCCGGAGAAAACCGCCTCGCGGTGATGGTGCTGCGCTGGAGTGACGGCAGTTATCTGGAAGATCAGGATATGTGGC
GGATGAGCGGCATTTTCCGTGACGTCTCGTTGCTGCATAAACCGACTACACAAATCAGCGATTTCCATGTTGCCACTCGCTT
TAATGATGATTTCAGCCGCGCTGTACTGGAGGCTGAAGTTCAGATGTGCGGCGAGTTGCGTGACTACCTACGGGTAACAGT
TTCTTTATGGCAGGGTGAAACGCAGGTCGCCAGCGGCACCGCGCCTTTCGGCGGTGAAATTATCGATGAGCGTGGTGGTTA
TGCCGATCGCGTCACACTACGTCTGAACGTCGAAAACCCGAAACTGTGGAGCGCCGAAATCCCGAATCTCTATCGTGCGGT
GGTTGAACTGCACACCGCCGACGGCACGCTGATTGAAGCAGAAGCCTGCGATGTCGGTTTCCGCGAGGTGCGGATTGAAA
ATGGTCTGCTGCTGCTGAACGGCAAGCCGTTGCTGATTCGAGGCGTTAACCGTCACGAGCATCATCCTCTGCATGGTCAGG
TCATGGATGAGCAGACGATGGTGCAGGATATCCTGCTGATGAAGCAGAACAACTTTAACGCCGTGCGCTGTTCGCATTATCC
GAACCATCCGCTGTGGTACACGCTGTGCGACCGCTACGGCCTGTATGTGGTGGATGAAGCCAATATTGAAACCCACGGCAT
GGTGCCAATGAATCGTCTGACCGATGATCCGCGCTGGCTACCGGCGATGAGCGAACGCGTAACGCGAATGGTGCAGCGCG
ATCGTAATCACCCGAGTGTGATCATCTGGTCGCTGGGGAATGAATCAGGCCACGGCGCTAATCACGACGCGCTGTATCGCT
GGATCAAATCTGTCGATCCTTCCCGCCCGGTGCAGTATGAAGGCGGCGGAGCCGACACCACGGCCACCGATATTATTTGCC
CGATGTACGCGCGCGTGGATGAAGACCAGCCCTTCCCGGCTGTGCCGAAATGGTCCATCAAAAAATGGCTTTCGCTACCTG
GAGAGACGCGCCCGCTGATCCTTTGCGAATACGCCCACGCGATGGGTAACAGTCTTGGCGGTTTCGCTAAATACTGGCAGG
CGTTTCGTCAGTATCCCCGTTTACAGGGCGGCTTCGTCTGGGACTGGGTGGATCAGTCGCTGATTAAATATGATGAAAACG
GCAACCCGTGGTCGGCTTACGGCGGTGATTTTGGCGATACGCCGAACGATCGCCAGTTCTGTATGAACGGTCTGGTCTTTG
CCGACCGCACGCCGCATCCAGCGCTGACGGAAGCAAAACACCAGCAGCAGTTTTTCCAGTTCCGTTTATCCGGGCAAACCA
TCGAAGTGACCAGCGAATACCTGTTCCGTCATAGCGATAACGAGCTCCTGCACTGGATGGTGGCGCTGGATGGTAAGCCGC
TGGCAAGCGGTGAAGTGCCTCTGGATGTCGCTCCACAAGGTAAACAGTTGATTGAACTGCCTGAACTACCGCAGCCGGAGA
GCGCCGGGCAACTCTGGCTCACAGTACGCGTAGTGCAACCGAACGCGACCGCATGGTCAGAAGCCGGACACATCAGCGCC
TGGCAGCAGTGGCGTCTGGCTGAAAACCTCAGCGTGACACTCCCCGCCGCGTCCCACGCCATCCCGCATCTGACCACCAG
CGAAATGGATTTTTGCATCGAGCTGGGTAATAAGCGTTGGCAATTTAACCGCCAGTCAGGCTTTCTTTCACAGATGTGGATT
GGCGATAAAAAACAACTGCTGACGCCGCTGCGCGATCAGTTCACCCGTGCACCGCTGGATAACGACATTGGCGTAAGTGAA
GCGACCCGCATTGACCCTAACGCCTGGGTCGAACGCTGGAAGGCGGCGGGCCATTACCAGGCCGAAGCAGCGTTGTTGCA
GTGCACGGCAGATACACTTGCTGATGCGGTGCTGATTACGACCGCTCACGCGTGGCAGCATCAGGGGAAAACCTTATTTAT
CAGCCGGAAAACCTACCGGATTGATGGTAGTGGTCAAATGGCGATTACCGTTGATGTTGAAGTGGCGAGCGATACACCGCA
TCCGGCGCGGATTGGCCTGAACTGCCAGCTGGCGCAGGTAGCAGAGCGGGTAAACTGGCTCGGATTAGGGCCGCAAGAA
AACTATCCCGACCGCCTTACTGCCGCCTGTTTTGACCGCTGGGATCTGCCATTGTCAGACATGTATACCCCGTACGTCTTCC
CGAGCGAAAACGGTCTGCGCTGCGGGACGCGCGAATTGAATTATGGCCCACACCAGTGGCGCGGCGACTTCCAGTTCAAC
ATCAGCCGCTACAGTCAACAGCAACTGATGGAAACCAGCCATCGCCATCTGCTGCACGCGGAAGAAGGCACATGGCTGAAT
ATCGACGGTTTCCATATGGGGATTGGTGGCGACGACTCCTGGAGCCCGTCAGTATCGGCGGAATTCCAGCTGAGCGCCGG
TCGCTACCATTACCAGTTGGTCTGGTGTCAAAAAtaagtcgaccggctgctaacaaagcccgaaaggaagctgagttggctgctgccaccgctgagcaataa
ctagcataaccccttggggcctctaaacgggtcttgaggggttttttg

| RNA *stx1 trigger* | RNA *stx1 trigger* |
|---|---|

*stx1 trigger*

ATAAATCGCCATTCGTTGACTACTTCTTATCTGGATTTAATGTCGCATAGTGGAACCTCACTGACGCAGTCTGTGGCAAGAGC
GATGTTACGGTTT

| $P_{T7}$-*stx1 switch*-*lacZ* | Plasmid encoding LacZ expression under T7 promoter and *stx1* toehold switch |
|---|---|

T7 Promoter-*stx1 switch*- *lacZ*-TrrnB-T7 Terminator

taatacgactcactatagggagaGGGCGTCAGTGAGGTTCCACTATGCGACATTAAATCCAGGGACTTTAGAACAGAGGAGATAAAGA
TGCTGGATTTAATTAACCTGGCGGCAGCGCAAAAGatgACCATGATTACGGATTCACTGGCCGTCGTTTTACAACGTCGTGAC
TGGGAAAACCCTGGCGTTACCCAACTTAATCGCCTTGCAGCACATCCCCCTTTCGCCAGCTGGCGTAATAGCGAAGAGGCC
CGCACCGATCGCCCTTCCCAACAGTTGCGCAGCCTGAATGGCGAATGGCGCTTTGCCTGGTTTCCGGCACCAGAAGCGGT
GCCGGAAAGCTGGCTGGAGTGCGATCTTCCTGAGGCCGATACTGTCGTCGTCCCCTCAAACTGGCAGATGCACGGTTACG
ATGCGCCCATCTACACCAACGTGACCTATCCCATTACGGTCAATCCGCCGTTTGTTCCCACGGAGAATCCGACGGGTTGTTA

CTCGCTCACATTTAATGTTGATGAAAGCTGGCTACAGGAAGGCCAGACGCGAATTATTTTTGATGGCGTTAACTCGGCGTTT
CATCTGTGGTGCAACGGGCGCTGGGTCGGTTACGGCCAGGACAGTCGTTTGCCGTCTGAATTTGACCTGAGCGCATTTTTA
CGCGCCGGAGAAAACCGCCTCGCGGTGATGGTGCTGCGCTGGAGTGACGGCAGTTATCTGGAAGATCAGGATATGTGGCG
GATGAGCGGCATTTTCCGTGACGTCTCGTTGCTGCATAAACCGACTACACAAATCAGCGATTTCCATGTTGCCACTCGCTTT
AATGATGATTTCAGCCGCGCTGTACTGGAGGCTGAAGTTCAGATGTGCGGCGAGTTGCGTGACTACCTACGGGTAACAGTT
TCTTTATGGCAGGGTGAAACGCAGGTCGCCAGCGGCACCGCGCCTTTCGGCGGTGAAATTATCGATGAGCGTGGTGGTTAT
GCCGATCGCGTCACACTACGTCTGAACGTCGAAAACCCGAAACTGTGGAGCGCCGAAATCCCGAATCTCTATCGTGCGGTG
GTTGAACTGCACACCGCCGACGGCACGCTGATTGAAGCAGAAGCCTGCGATGTCGGTTTCCGCGAGGTGCGGATTGAAAA
TGGTCTGCTGCTGCTGAACGGCAAGCCGTTGCTGATTCGAGGCGTTAACCGTCACGAGCATCATCCTCTGCATGGTCAGGT
CATGGATGAGCAGACGATGGTGCAGGATATCCTGCTGATGAAGCAGAACAACTTTAACGCCGTGCGCTGTTCGCATTATCC
GAACCATCCGCTGTGGTACACGCTGTGCGACCGCTACGGCCTGTATGTGGTGGATGAAGCCAATATTGAAACCCACGGCAT
GGTGCCAATGAATCGTCTGACCGATGATCCGCGCTGGCTACCGGCGATGAGCGAACGCGTAACGCGAATGGTGCAGCGCG
ATCGTAATCACCCGAGTGTGATCATCTGGTCGCTGGGGAATGAATCAGGCCACGGCGCTAATCACGACGCGCTGTATCGCT
GGATCAAATCTGTCGATCCTTCCCGCCCGGTGCAGTATGAAGGCGGCGGAGCCGACACCACGGCCACCGATATTATTTGCC
CGATGTACGCGCGCGTGGATGAAGACCAGCCCTTCCCGGCTGTGCCGAAATGGTCCATCAAAAAATGGCTTTCGCTACCTG
GAGAGACGCGCCCGCTGATCCTTTGCGAATACGCCCACGCGATGGGTAACAGTCTTGGCGGTTTCGCTAAATACTGGCAGG
CGTTTCGTCAGTATCCCCGTTTACAGGGCGGCTTCGTCTGGGACTGGGTGGATCAGTCGCTGATTAAATATGATGAAAACG
GCAACCCGTGGTCGGCTTACGGCGGTGATTTTGGCGATACGCCGAACGATCGCCAGTTCTGTATGAACGGTCTGGTCTTTG
CCGACCGCACGCCGCATCCAGCGCTGACGGAAGCAAAACACCAGCAGCAGTTTTTCCAGTTCCGTTTATCCGGGCAAACCA
TCGAAGTGACCAGCGAATACCTGTTCCGTCATAGCGATAACGAGCTCCTGCACTGGATGGTGGCGCTGGATGGTAAGCCGC
TGGCAAGCGGTGAAGTGCCTCTGGATGTCGCTCCACAAGGTAAACAGTTGATTGAACTGCCTGAACTACCGCAGCCGGAGA
GCGCCGGGCAACTCTGGCTCACAGTACGCGTAGTGCAACCGAACGCGACCGCATGGTCAGAAGCCGGACACATCAGCGCC
TGGCAGCAGTGGCGTCTGGCTGAAAACCTCAGCGTGACACTCCCCGCCGCGTCCCACGCCATCCCGCATCTGACCACCAG
CGAAATGGATTTTTGCATCGAGCTGGGTAATAAGCGTTGGCAATTTAACCGCCAGTCAGGCTTTCTTTCACAGATGTGGATT
GGCGATAAAAAACAACTGCTGACGCCGCTGCGCGATCAGTTCACCCGTGCACCGCTGGATAACGACATTGGCGTAAGTGAA
GCGACCCGCATTGACCCTAACGCCTGGGTCGAACGCTGGAAGGCGGCGGGCCATTACCAGGCCGAAGCAGCGTTGTTGCA
GTGCACGGCAGATACACTTGCTGATGCGGTGCTGATTACGACCGCTCACGCGTGGCAGCATCAGGGGAAAACCTTATTTAT
CAGCCGGAAAACCTACCGGATTGATGGTAGTGGTCAAATGGCGATTACCGTTGATGTTGAAGTGGCGAGCGATACACCGCA
TCCGGCGCGGATTGGCCTGAACTGCCAGCTGGCGCAGGTAGCAGAGCGGGTAAACTGGCTCGGATTAGGGCCGCAAGAA
AACTATCCCGACCGCCTTACTGCCGCCTGTTTTGACCGCTGGGATCTGCCATTGTCAGACATGTATACCCCGTACGTCTTCC
CGAGCGAAAACGGTCTGCGCTGCGGGACGCGCGAATTGAATTATGGCCCACACCAGTGGCGCGGCGACTTCCAGTTCAAC
ATCAGCCGCTACAGTCAACAGCAACTGATGGAAACCAGCCATCGCCATCTGCTGCACGCGGAAGAAGGCACATGGCTGAAT
ATCGACGGTTTCCATATGGGGATTGGTGGCGACGACTCCTGGAGCCCGTCAGTATCGGCGGAATTCCAGCTGAGCGCCGG
TCGCTACCATTACCAGTTGGTCTGGTGTCAAAAAtaataaggatctgaagcttgggcccgaacaaaaactcatctcagaagaggatctgaatagcgccgtcg
accatcatcatcatcatcattgagtttaaacggtctccagcttggctgttttggcggatgagagaagattttcagcctgatacagattaaatcagaacgcagaagcggtctgataaaacag
aatttgcctggcggcagtagcgcggtggtcccacctgaccccatgccgaactcagaagtgaaacgccgtagcgccgatggtagtgtggggtctccccatgcgagagtagggaactg
ccaggcatcaaataaaacgaaaggctcagtcgaaagactgggcctttcgttttatctgttgtttgtcggtgaactggatcgtcgaccggctgctaacaaagcccgaaaggaagctgagt
tggctgctgccaccgctgagcaataacctagcataaccccttggggcctctaaacgggtcttgaggggttttttg

| **P$_{T7}$-*stx2 trigger*** | Linear DNA encoding expression of RNA *stx2* trigger under a T7 Promoter. Additional nucleotides (~30bp) were included in 5’ and 3’ end as extra protection against nuclease degradation. |
|---|---|

T7 Promoter-*stx2 trigger*

ggaaaaacgccagcaacgcgatcccgcgaaattaatacgactcactataggGTATCCTATTCCCGGGAGTTTACGATAGACTTTTCGACCCAACAA
AGTTATGTCTCTTCGTTAAATAGTATACGGACAGAGATATCGACCCCTCTTGAACATATATCaaaaaaaacgccgcctttcggcggcgtttg

| **P$_{T7}$-*stx2 switch-lacZ*** | Plasmid encoding LacZ expression under T7 promoter and *stx2* toehold switch |
|---|---|

T7 Promoter-*stx2 switch*- *lacZ*- TrrnB-T7 Terminator

taatacgactcactatagggagaGGGATACTATTTAACGAAGAGACATAACTTTGTTGGGTCGGACTTTAGAACAGAGGAGATAAAGAT
GGACCCAACAAAGatgACCATGATTACGGATTCACTGGCCGTCGTTTTACAACGTCGTGACTGGGAAAACCCTGGCGTTACC
CAACTTAATCGCCTTGCAGCACATCCCCCTTTCGCCAGCTGGCGTAATAGCGAAGAGGCCCGCACCGATCGCCCTTCCCAA
CAGTTGCGCAGCCTGAATGGCGAATGGCGCTTTGCCTGGTTTCCGGCACCAGAAGCGGTGCCGGAAAGCTGGCTGGAGTG
CGATCTTCCTGAGGCCGATACTGTCGTCGTCCCCTCAAACTGGCAGATGCACGGTTACGATGCGCCCATCTACACCAACGT
GACCTATCCCATTACGGTCAATCCGCCGTTTGTTCCCACGGAGAATCCGACGGGTTGTTACTCGCTCACATTTAATGTTGAT
GAAAGCTGGCTACAGGAAGGCCAGACGCGAATTATTTTTGATGGCGTTAACTCGGCGTTTCATCTGTGGTGCAACGGGCGC
TGGGTCGGTTACGGCCAGGACAGTCGTTTGCCGTCTGAATTTGACCTGAGCGCATTTTTACGCGCCGGAGAAAACCGCCTC
GCGGTGATGGTGCTGCGCTGGAGTGACGGCAGTTATCTGGAAGATCAGGATATGTGGCGGATGAGCGGCATTTTCCGTGA
CGTCTCGTTGCTGCATAAACCGACTACACAAATCAGCGATTTCCATGTTGCCACTCGCTTTAATGATGATTTCAGCCGCGCT

GTACTGGAGGCTGAAGTTCAGATGTGCGGCGAGTTGCGTGACTACCTACGGGTAACAGTTTCTTTATGGCAGGGTGAAACG
CAGGTCGCCAGCGGCACCGCGCCTTTCGGCGGTGAAATTATCGATGAGCGTGGTGGTTATGCCGATCGCGTCACACTACG
TCTGAACGTCGAAAACCCGAAACTGTGGAGCGCCGAAATCCCGAATCTCTATCGTGCGGTGGTTGAACTGCACACCGCCGA
CGGCACGCTGATTGAAGCAGAAGCCTGCGATGTCGGTTTCCGCGAGGTGCGGATTGAAAATGGTCTGCTGCTGCTGAACG
GCAAGCCGTTGCTGATTCGAGGCGTTAACCGTCACGAGCATCATCCTCTGCATGGTCAGGTCATGGATGAGCAGACGATGG
TGCAGGATATCCTGCTGATGAAGCAGAACAACTTTAACGCCGTGCGCTGTTCGCATTATCCGAACCATCCGCTGTGGTACAC
GCTGTGCGACCGCTACGGCCTGTATGTGGTGGATGAAGCCAATATTGAAACCCACGGCATGGTGCCAATGAATCGTCTGAC
CGATGATCCGCGCTGGCTACCGGCGATGAGCGAACGCGTAACGCGAATGGTGCAGCGCGATCGTAATCACCCGAGTGTGA
TCATCTGGTCGCTGGGGAATGAATCAGGCCACGGCGCTAATCACGACGCGCTGTATCGCTGGATCAAATCTGTCGATCCTT
CCCGCCCGGTGCAGTATGAAGGCGGCGGAGCCGACACCACGGCCACCGATATTATTTGCCCGATGTACGCGCGCGTGGAT
GAAGACCAGCCCTTCCCGGCTGTGCCGAAATGGTCCATCAAAAAATGGCTTTCGCTACCTGGAGAGACGCGCCCGCTGATC
CTTTGCGAATACGCCCACGCGATGGGTAACAGTCTTGGCGGTTTCGCTAAATACTGGCAGGCGTTTCGTCAGTATCCCCGTT
TACAGGGCGGCTTCGTCTGGGACTGGGTGGATCAGTCGCTGATTAAATATGATGAAAACGGCAACCCGTGGTCGGCTTACG
GCGGTGATTTTGGCGATACGCCGAACGATCGCCAGTTCTGTATGAACGGTCTGGTCTTTGCCGACCGCACGCCGCATCCAG
CGCTGACGGAAGCAAAACACCAGCAGCAGTTTTTCCAGTTCCGTTTATCCGGGCAAACCATCGAAGTGACCAGCGAATACC
TGTTCCGTCATAGCGATAACGAGCTCCTGCACTGGATGGTGGCGCTGGATGGTAAGCCGCTGGCAAGCGGTGAAGTGCCT
CTGGATGTCGCTCCACAAGGTAAACAGTTGATTGAACTGCCTGAACTACCGCAGCCGGAGAGCGCCGGGCAACTCTGGCTC
ACAGTACGCGTAGTGCAACCGAACGCGACCGCATGGTCAGAAGCCGGACACATCAGCGCCTGGCAGCAGTGGCGTCTGGC
TGAAAACCTCAGCGTGACACTCCCCGCCGCGTCCCACGCCATCCCGCATCTGACCACCAGCGAAATGGATTTTTGCATCGA
GCTGGGTAATAAGCGTTGGCAATTTAACCGCCAGTCAGGCTTTCTTTCACAGATGTGGATTGGCGATAAAAAACAACTGCTG
ACGCCGCTGCGCGATCAGTTCACCCGTGCACCGCTGGATAACGACATTGGCGTAAGTGAAGCGACCCGCATTGACCCTAAC
GCCTGGGTCGAACGCTGGAAGGCGGCGGGCCATTACCAGGCCGAAGCAGCGTTGTTGCAGTGCACGGCAGATACACTTGC
TGATGCGGTGCTGATTACGACCGCTCACGCGTGGCAGCATCAGGGGAAAACCTTATTTATCAGCCGGAAAACCTACCGGAT
TGATGGTAGTGGTCAAATGGCGATTACCGTTGATGTTGAAGTGGCGAGCGATACACCGCATCCGGCGCGGATTGGCCTGAA
CTGCCAGCTGGCGCAGGTAGCAGAGCGGGTAAACTGGCTCGGATTAGGGCCGCAAGAAAACTATCCCGACCGCCTTACTG
CCGCCTGTTTTGACCGCTGGGATCTGCCATTGTCAGACATGTATACCCCGTACGTCTTCCCGAGCGAAAACGGTCTGCGCT
GCGGGACGCGCGAATTGAATTATGGCCCACACCAGTGGCGCGGCGACTTCCAGTTCAACATCAGCCGCTACAGTCAACAG
CAACTGATGGAAACCAGCCATCGCCATCTGCTGCACGCGGAAGAAGGCACATGGCTGAATATCGACGGTTTCCATATGGGG
ATTGGTGGCGACGACTCCTGGAGCCCGTCAGTATCGGCGGAATTCCAGCTGAGCGCCGGTCGCTACCATTACCAGTTGGT
CTGGTGTCAAAAAtaataaggatctgaagcttgggcccgaacaaaaactcatctcagaagaggatctgaatagcgccgtcgaccatcatcatcatcatcattgagtttaaacg
gtctccagcttggctgttttggcggatgagagaagattttcagcctgatacagattaaatcagaacgcagaagcggtctgataaaacagaatttgcctggcggcagtagcgcggtggtc
ccacctgaccccatgccgaactcagaagtgaaacgccgtagcgccgatggtagtgtggggtctccccatgcgagagtagggaactgccaggcatcaaataaaacgaaaggctca
gtcgaaagactgggcctttcgttttatctgttgtttgtcggtgaactggatcgtcgaccggctgctaacaaagcccgaaaggaagctgagttggctgctgccaccgctgagcaataaccta
gcataaccccttggggcctctaaacgggtcttgaggggttttttg

| **$P_{T7}$-*gfp*** | Plasmid encoding super folder GFP expression under T7 promoter with a strong ribosomal binding site |
|---|---|

T7 Promoter-Stability Hairpin-StrongRBS-*gfp*-T7 Terminator

taatacgactcactatagggagaccacaacggtttccctctagaaataattttgtttaactttaagaaggagatatacatATGAGCAAAGGTGAAGAACTGTTTACCG
GCGTTGTGCCGATTCTGGTGGAACTGGATGGCGATGTGAACGGTCACAAATTCAGCGTGCGTGGTGAAGGTGAAGGCGAT
GCCACGATTGGCAAACTGACGCTGAAATTTATCTGCACCACCGGCAAACTGCCGGTGCCGTGGCCGACGCTGGTGACCAC
CCTGACCTATGGCGTTCAGTGTTTTAGTCGCTATCCGGATCACATGAAACGTCACGATTTCTTTAAATCTGCAATGCCGGAAG
GCTATGTGCAGGAACGTACGATTAGCTTTAAAGATGATGGCAAATATAAAACGCGCGCCGTTGTGAAATTTGAAGGCGATAC
CCTGGTGAACCGCATTGAACTGAAAGGCACGGATTTTAAAGAAGATGGCAATATCCTGGGCCATAAACTGGAATACAACTTT
AATAGCCATAATGTTTATATTACGGCGGATAAACAGAAAAATGGCATCAAAGCGAATTTTACCGTTCGCCATAACGTTGAAGA
TGGCAGTGTGCAGCTGGCAGATCATTATCAGCAGAATACCCCGATTGGTGATGGTCCGGTGCTGCTGCCGGATAATCATTAT
CTGAGCACGCAGACCGTTCTGTCTAAAGATCCGAACGAAAAAGGCACGCGGGACCACATGGTTCTGCACGAATATGTGAAT
GCGGCAGGTATTACGTGGAGCCATCCGCAGTTCGAAAAATAAgtcgaccggctgctaacaaagcccgaaaggaagctgagttggctgctgccaccgct
gagcaataactagcataaccccttggggcctctaaacgggtcttgaggggttttttg

| **$P_{T7}$-MGA** | Plasmid encoding malachite green RNA aptamer expression under the T7 promoter |
|---|---|

T7 Promoter-MGA-T7 Terminator

taatacgactcactatagggGGGATCCCGACTGGCGAGAGCCAGGTAACGAATGGATCGGGTCGGCATGGCATCTCCACCTCCTCG
CGGTCCGACCTGGGCATCCGAAGGAGGACGTCGTCCACTCGGATGGCTAAGGGAGAGCTCGGATCCGGCTGCTAACAAAG
CCCGAAAGGAAGCTGAGTTGGCTGCTGCCACCGCTGAGCAATAACTAGCATAAtagcataaccccttggggcctctaaacgggtcttgaggggtt
ttttg

**Table S2.** Description of lysate, plasmid concentrations, and reaction additives present in CFE reactions in each figure.

| Figures | Detecting | Plasmids/ RNA Transcripts | Reaction Additives |
|---|---|---|---|
| 1A | N/A | N/A | 0-25 mM glucose |
| 1B | N/A | 0-2 nM $P_{T7}$-*lacZ* | |
| 2B | Zinc in water | 1 nM $P_{T7}$-*zntR*<br>2 nM $P_{ZntA}$-*lacZ* | |
| 2C | Zinc in serum | 2 nM $P_{T7}$-*zntR*<br>8 nM $P_{ZntA}$-*lacZ* | 1.5 % RNase Inhibitor Murine<br>25 % Pooled human serum |
| 3B | RNA *stx1 trigger* | 4 nM $P_{T7}$- *stx1 switch-lacZ* | 0.5 % RNase Inhibitor Murine |
| 3C | Linear DNA *stx2 trigger* | 5 nM $P_{T7}$- *stx2 switch-lacZ* | 4 $\mu$M GamS Protein |
| S1A | N/A | 1 nM $P_{T7}$-*gfp* | -/+ 10 mM Naproxen sodium |
| S1B | N/A | N/A | 5% BL21 Star (DE3) lysate containing LacZ<br>0.6 mg/mL CPRG |
| S1C | N/A | 5 nM $P_{T7}$-MGA | 1 mM Malachite Green dye<br>-/+ 10 mM Naproxen sodium |
| S1D | N/A | 0.5 $\mu$M *gfp* RNA | -/+ 10 mM Naproxen sodium |
| S1E | N/A | N/A | 10 mM Naproxen sodium<br>0-25 mM glucose |
| S2B | N/A | N/A | 25% Pooled human serum with glucose dosed in |
| S2C | N/A | 1 nM $P_{T7}$-*gfp* | 1.5 % RNase Inhibitor Murine<br>25% Pooled human serum with glucose dosed in |
| S3B, C | Linear DNA *stx2 trigger* | 5 nM $P_{T7}$-*stx2 switch-lacZ* | 4 $\mu$M GamS Protein |

**Table S3:** Primers used for trigger DNA amplification from *E. coli* O157: H7 genomic DNA template. Lowercase, unlabeled sequences are protective regions to decrease endonuclease degradation. Highlighted sequences indicate the T7 promoter, and uppercase sequences are primer annealing regions. Target-specific primers were designed to bind at least 20 base pairs before and after the actual trigger sequence to prevent unintended primer activation of switches.

| Amplifying | Primer Sequence |
|---|---|
| *E. coli* | Fwd:<br>ggaaaaacgccagcaacgcgatcccgcgaaattaatacgactcactataggCAAACTACGACGTCATCATTTAGC<br><br>Rev:<br>caaacgccgccgaaaggcggcgttttttttGTTGACGGTTCAAACGTGGAAA<br><br>Genomic Target: AraC family transcriptional regulator.<br>Accession # LR881938. Region: 315725...315888.<br>TCAAACTACGACGTCATCATTTAGCCAGATGTATGAAGAATTTTAAGACGGATATCTATTTCGTTTCCAC<br>GTTTGAACCGTCAACAAAATCGGTCGATTTGCTCACGGTTGAAACTTTTGCTGGTACGGTATGTGAATAT<br>GCTGACATGCCAAAAGAGTGGACA |
| *stx1* | Fwd:<br>ggaaaaacgccagcaacgcgatcccgcgaaattaatacgactcactataggATAAATCGCCATTCGTTGACTACT<br><br>Rev:<br>caaacgccgccgaaaggcggcgttttttttAAACCGTAACATCGCTCTTGCCA<br><br>Genomic Target: Shiga Toxin 1.<br>Accession #: BA000007. Region: 2924904...2925851.<br>ATGAAAATAATTATTTTTAGAGTGCTAACTTTTTTCTTTGTTATCTTTTCAGTTAATGTGGTTGCGAAGGAA<br>TTTACCTTAGACTTCTCGACTGCAAAGACGTATGTAGATTCGCTGAATGTCATTCGCTCTGCAATAGGTA<br>CTCCATTACAGACTATTTCATCAGGAGGTACGTCTTTACTGATGATTGATAGTGGCACAGGGGATAATTT<br>GTTTGCAGTTGATGTCAGAGGGATAGATCCAGAGGAAGGGCGGTTTAATAATCTACGGCTTATTGTTGA<br>ACGAAATAATTTATATGTGACAGGATTTGTTAACAGGACAAATAATGTTTTTTATCGCTTTGCTGATTTTTC<br>ACATGTTACCTTTCCAGGTACAACAGCGGTTACATTGTCTGGTGACAGTAGCTATACCACGTTACAGCG<br>TGTTGCAGGGATCAGTCGTACGGGGATGCAGATAAATCGCCATTCGTTGACTACTTCTTATCTGGATTTA<br>ATGTCGCATAGTGGAACCTCACTGACGCAGTCTGTGGCAAGAGCGATGTTACGGTTTGTTACTGTGACA<br>GCTGAAGCTTTACGTTTTCGGCAAATACAGAGGGGATTTCGTACAACACTGGATGATCTCAGTGGGCGT<br>TCTTATGTAATGACTGCTGAAGATGTTGATCTTACATTGAACTGGGGAAGGTTGAGTAGTGTCCTGCCTG<br>ATTATCATGGACAAGACTCTGTTCGTGTAGGAAGAATTTCTTTTGGAAGCATTAATGCAATTCTGGGAAG<br>CGTGGCATTAATACTGAATTGTCATCATCATGCATCGCGAGTTGCCAGAATGGCATCTGATGAGTTTCCT<br>TCTATGTGTCCGGCAGATGGAAGAGTCCGTGGGATTACGCACAATAAAATATTGTGGGATTCATCCACT<br>CTGGGGGCAATTCTGATGCGCAGAACTATTAGCAGTTG |
| *stx2* | Fwd:<br>ggaaaaacgccagcaacgcgatcccgcgaaattaatacgactcactataggGTATCCTATTCCCGGGAGTTTACGATAGACTT<br>TTC<br><br>Rev:<br>caaacgccgccgaaaggcggcgttttttttGATATATGTTCAAGAGGGGTCGATATCTCTGTCCG<br><br>Genomic Target: Shiga Toxin 2.<br>Accession #: BA000007. Region: 1267107…1268066.<br><br>ATGAAGTGTATATTATTTAAATGGGTACTGTGCCTGTTACTGGGTTTTTCTTCGGTATCCTATTCCCGGG<br>AGTTTACGATAGACTTTTCGACCCAACAAAGTTATGTCTCTTCGTTAAATAGTATACGGACAGAGATATC<br>GACCCCTCTTGAACATATATCTCAGGGGACCACATCGGTGTCTGTTATTAACCACACCCCACCGGGCAG<br>TTATTTTGCTGTGGATATACGAGGGCTTGATGTCTATCAGGCGCGTTTTGACCATCTTCGTCTGATTATT<br>GAGCAAAATAATTTATATGTGGCCGGGTTCGTTAATACGGCAACAAATACTTTCTACCGTTTTTCAGATTT<br>TACACATATATCAGTGCCCGGTGTGACAACGGTTTCCATGACAACGGACAGCAGTTATACCACTCTGCA |

| | |
|---|---|
| | ACGTGTCGCAGCGCTGGAACGTTCCGGAATGCAAATCAGTCGTCACTCACTGGTTTCATCATATCTGGC<br>GTTAATGGAGTTCAGTGGTAATACAATGACCAGAGATGCATCCAGAGCAGTTCTGCGTTTTGTCACTGT<br>CACAGCAGAAGCCTTACGCTTCAGGCAGATACAGAGAGAATTTCGTCAGGCACTGTCTGAAACTGCTCC<br>TGTGTATACGATGACGCCGGGAGACGTGGACCTCACTCTGAACTGGGGGCGAATCAGCAATGTGCTTC<br>CGGAGTATCGGGGAGAGGATGGTGTCAGAGTGGGGAGAATATCCTTTAATAATATATCAGCGATACTG<br>GGGACTGTGGCCGTTATACTGAATTGCCATCATCAGGGGGCGCGTTCTGTTCGCGCCGTGAATGAAGA<br>GAGTCAACCAGAATGTCAGATAACTGGCGACAGGCCTGTTATAAAAATAAACAATACATTATGGGAAAGT<br>AATACAGCTGCAGCGTTTCTGAACAGAAAGTCACAGTTTTTATATACAACGGGTAAATAA |

**References:**

[1] Sun, Z. Z., Hayes, C. A., Shin, J., Caschera, F., Murray, R. M., and Noireaux, V. (2013) Protocols for implementing an Escherichia coli based TX-TL cell-free expression system for synthetic biology, *J Vis Exp*, e50762.

[2] Takahashi, M. K., Tan, X., Dy, A. J., Braff, D., Akana, R. T., Furuta, Y., Donghia, N., Ananthakrishnan, A., and Collins, J. J. (2018) A low-cost paper-based synthetic biology platform for analyzing gut microbiota and host biomarkers, *Nat Commun 9*, 3347.
